# Loss of epigenetic regulation disrupts lineage integrity, reactivates multipotency and promotes breast cancer

**DOI:** 10.1101/2021.10.22.465428

**Authors:** Ellen R. Langille, Khalid N. Al-Zahrani, Zhibo Ma, Ahmad Malik, Sampath K. Loganathan, Dan Trcka, Jeff Liu, Katelyn Kozma, Ricky Tsai, Katie Teng, Roderic Espin, Seda Barutcu, Thomas Nguyen, Rod Bremner, Hartland W. Jackson, Erik S. Knudsen, Gary Bader, Sean E. Egan, Miquel Angel Pujana, Jeff Wrana, Geoffrey M. Wahl, Daniel Schramek

## Abstract

Systematically investigating the scores of genes mutated in cancer and discerning real drivers from inconsequential bystanders is a prerequisite for Precision Medicine, but remains challenging. Here, we developed a somatic CRISPR/Cas9 mutagenesis screen to study 215 recurrent ‘long-tail’ breast cancer genes, which revealed epigenetic regulation as major tumor suppressive mechanism. We report that core or accessory components of the COMPASS histone methylase complex including *KMT2C*, *KDM6A*, *BAP1* and *ASXL2* (“EpiDrivers”) cooperate with *PIK3CA*^H1047R^ to transform mouse and human breast epithelial cells. Mechanistically, we find that Cre-mediated activation of *PIK3CA*^H1047R^ elicited an aberrant alveolar lactation program in luminal cells, which was exacerbated upon loss of EpiDrivers. Remarkably, EpiDriver loss in basal cells also triggered an alveolar-like lineage conversion and accelerated formation of luminal-like tumors, suggesting a basal origin for luminal tumors. As EpiDrivers are mutated in 39% of human breast cancers, lineage infidelity and lactation mimicry may significantly contribute to early steps of breast cancer progression.

**Statement of significance:** Infrequently mutated genes comprise most of the mutational burden in breast tumors but are poorly understood. *In-vivo* CRISPR screening identified functional tumor suppressors that converged on epigenetic regulation. Loss of epigenetic regulators accelerated tumorigenesis and revealed lineage infidelity and aberrant expression of lactation genes as potential early events in tumorigenesis.

## Introduction

New genomic technologies hold the promise of revolutionizing cancer therapy by allowing treatment decisions guided by a tumor’s genetic make-up. However, converting genetic discoveries into tangible clinical benefits requires a deeper understanding of the molecular and cellular mechanisms that underlie disease progression. In breast cancer, only a few genes such as *TP53* and *PIK3CA* are mutated at high frequencies (∼30-50%), while the vast majority are mutated at low frequencies comprising a ‘long-tail’ distribution (1-3). Whether these long-tail genes functionally contribute to breast cancer progression constitutes a significant knowledge gap. Although these mutations seem to be under positive selection, they are only found in a relatively small subset of patients. It has been proposed that these infrequently mutated genes may individually confer small fitness advantages but when combined might synergize to increase fitness (additive-effects model) (4-6). Alternatively, long-tail mutations may be sufficient to promote tumorigenesis but are rarely mutated, either because they work in different ways to produce the same phenotype (phenotypic convergence), or they affect the same pathway or molecular mechanism (pathway convergence)(7). Recently, we exposed the latter mechanism in in head and neck cancer, where rarely mutated long tail genes converge to inactivate NOTCH signaling (8). Here, we report an *in vivo* CRISPR/Cas9 screening strategy to identify which long-tail breast cancer genes and associated molecular pathways cooperate with the oncogenic *PIK3CA^H1047R^* mutation to accelerate breast cancer progression.

We tested 215 long-tail genes and identified several functionally important breast cancer genes, many of which converge on regulating histone modifications and enhancer activity (EpiDrivers). Single-cell multi-omics profiling of EpiDrivers-mutant mammary glands reveals increased cell state plasticity and lactation mimicry associated with an aberrant alveolar differentiation program during the early specification of luminal breast cancer. Interestingly, EpiDrivers loss in basal cells triggers basal-to-alveolar linage conversion and accelerated tumor formation. Importantly, EpiDriver mutations are found in ∼39% of breast cancer patients, highlighting the notion that different genes converge to produce the same cell plasticity to facilitates cancer development.

## RESULTS

### Direct *in vivo* CRISPR gene editing in the Mouse Mammary Gland

First, we developed a multiplexed CRISPR/Cas9 knock-out approach in the mammary gland of tumor-prone mice. As *PIK3CA* is the most commonly mutated oncogene in breast cancer, we crossed conditional Lox-Stop-Lox-(LSL)-*Pik3ca^H1047R^* mice to LSL-*Cas9-GFP* transgenic mice (termed *Pik3ca^H1047R^*;*Cas9* mice). Intraductal microinjections of a lentivirus that expresses an sgRNA and Cre recombinase (Lenti-sgRNA-Cre) led to excision of Lox-Stop-Lox cassettes and expression of *Cas9*, *GFP* and oncogenic *Pik3ca^H1047R^* in the mammary epithelium (**Fig. 1A**). To validate the efficacy of CRISPR/Cas9-mediated mutagenesis, we injected sgRNAs targeting *GFP* or the heme biosynthesis gene *Urod.* Knock-out of *GFP* was detected as a 86±6% reduction in green fluorescence in transduced cells, whereas knock-out of *Urod* was detected as an accumulation of unprocessed fluorescent porphyrins in 30%±8% of cells (**Supplementary Fig. S1A-D**) (9). Moreover, *Pik3ca^H1047R^*;*Cas9* mice transduced with an sgRNA targeting *Trp53* developed tumors much faster than littermate mice transduced with a control sgRNA targeting the permissive Tigre locus (median tumor-free survival of 83 versus 152 days) (**Supplementary Fig. S1E**). Together, these data demonstrate that this approach recapitulates cooperation between oncogenic *Pik3ca* and *Trp53* loss-of-function (10,11) and can be used to test for genetic interaction between breast cancer genes.

**Fig. 1.**
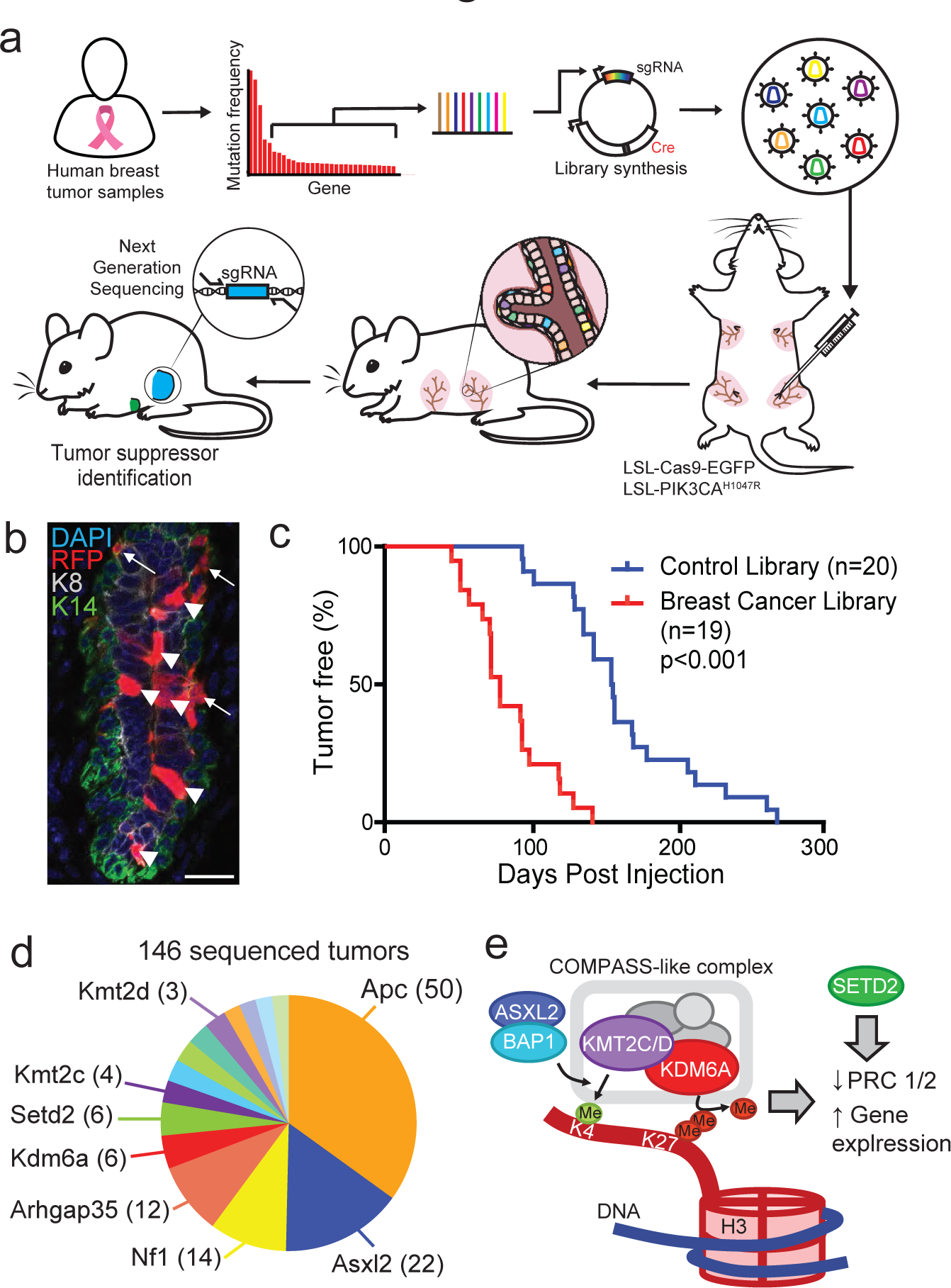
In vivo CRISPR screen reveals novel epigenetic breast cancer tumors suppressors ‘EpiDrivers’. **a**, Experimental design for in vivo CRISPR screen, showing gene selection from long-tail mutations, intraductal injection of lentiviral libraries and tumor sequencing. **b**, Mammary epithelium transduced with lentiviral RFP. Arrows denote basal cells and arrow heads denote luminal cells. Scale bar = 25µm. **c**, Tumor-free survival of Pik3ca^H1047R^;Cas9 mice transduced with a sgRNA library targeting putative breast cancer genes or a control sgRNA library. **d**, Pie chart showing putative tumor suppressor genes with enriched sgRNAs in tumor DNA obtained (number of tumors are denoted in brackets). **e**, Schematic of COMPASS-like and ASXL/BAP1 complexes on epigenetic control of gene expression.

### CRISPR Screen Identifies Histone Modifiers as Breast Cancer Driver Genes

In breast cancer, 215 long-tail genes show somatic mutations in 2-20% of patients (10,12). To assess these genes *in vivo*, we established a sgRNA library targeting the corresponding mouse orthologs (4 sgRNAs/gene; 860 sgRNAs) as well as a library of 420 non-targeting control sgRNAs (**Supplementary Table S1**).

Next, we optimized the parameters for an *in vivo* CRISPR screen. Using a mixture of lentiviral Lenti-GFP and Lenti-RFP we determined the viral titer that transduces the mammary epithelium at clonal density (MOI<1). Higher viral titers were associated with double infections, whereas a 15% overall transduction level minimized double infections while generating sufficient clones to screen (**Supplementary Fig. S1F and S1G**). Flow cytometry revealed that the third and fourth mammary gland each contain >3.5x10^5^ epithelial cells, and that EPCAM^hi^/CD49f^mid^ luminal cells showed a higher infectivity (∼30%) compared to basal EPCAM^mid^/CD49f^hi^ cells (∼5%) (**Fig. 1B** and **Supplementary Fig. S1H and S1I**). Thus, at a transduction level of 15% and a pool of 860 sgRNAs, each sgRNA would be introduced into an average of 60 individual cells within a single gland.

To uncover long-tail genes that cooperate with oncogenic PI3K signaling, we introduced the viral libraries into the third and fourth pairs of mammary glands of 19 *Pik3ca^H1047R^*;*Cas9* mice, resulting in an overall coverage of >4000 clones per sgRNA. Next generation sequencing confirmed efficient lentiviral transduction of all sgRNAs (**Supplementary Fig. S2A**). Importantly, *Pik3ca^H1047R^;Cas9* female mice transduced with the long-tail breast cancer sgRNA library developed mammary tumors significantly faster than littermates transduced with the control sgRNA library (74 versus 154 days; p<0.0001) (**Fig. 1C**). This result was reminiscent of the effect seen upon mutation of *Trp53* (**Supplementary Fig. S1E**), indicating the existence of strong tumor suppressors within the long-tail of breast cancer-associated genes.

We examined the sgRNA representation in 146 tumors to determine the targets responsible for accelerating mammary tumor formation. Most tumors showed strong enrichment for a single or sometimes two sgRNAs (**Supplementary Fig. S2B**). We prioritized genes that were targeted by ≥2 sgRNAs and knocked-out in multiple tumors, resulting in 29 candidate tumor suppressor genes (**Supplementary Table S2**). These candidates included well-known tumor suppressors, such as *Apc* or *Nf1*, as well as genes with poorly understood function, such as *Arhgap35.* Intriguingly, several genes encoded histone and DNA modifying enzymes, such as *Arid5b*, *Asxl2*, *Kdm6a* (*Utx*), *Kmt2a* (*Mll1*), *Kmt2c* (*Mll3*), and *Kmt2d* (*Mll4*), indicating a convergence on epigenetic regulation (**Fig. 1D, Supplementary Fig. S2C**).

### *Kdm6a, Kmt2c, Asxl2, Bap1, Setd2* and *Apc* suppress breast cancer in mice

*KMT2C* and *KMT2D* encode partly redundant histone methyltransferases within the ‘complex of proteins associated with SET1’ (COMPASS)-like complex, which also contains the histone demethylase KDM6A. The KMT2C/D-COMPASS-like complex catalyzes the mono-methylation of lysine 4 as well as demethylation of lysine 27 in histone H3 (H3K4me1/H3K27) at distal enhancers, facilitating recruitment of the CBP/p300 H3K27 histone acetylase (HAT), which ultimately primes enhancers for gene activation (13,14). The KMT2C/D-COMPASS-like complex is recruited to enhancers by the BAP1-ASXL1/2 complex, which facilitates enhancer priming (15,16). In addition, the methyltransferase SETD2 deposits H3K36me3 marks at active enhancers and transcribed gene bodies (17,18). Thus, our top hits converge on regulating enhancer function (**Fig. 1E**).

We validated each hit by injecting *Pik3ca^H1047R^; Cas9* mice individually with one sgRNA from the library and one newly designed sgRNA targeting *Asxl2*, *Kdm6a*, *Kmt2c*, and *Setd2* (termed EpiDrivers) or *Trp53* and *Apc* as controls. We also transduced mice with sgRNAs targeting *Bap1*, which was not in the original library. All transduced mice developed multiple highly proliferative breast tumors with much shorter latencies compared to mice transduced with non-targeting control sgRNAs (sgNT) (**Fig. 2A**; **Supplementary Fig. S2D and S2E**). All tested tumors harbored bi-allelic frame-shift mutations in the target genes, and western blot analysis confirmed loss of Apc, Asxl2, Kdm6a, and p53 expression (**Supplementary Fig. S2F-K)**.

**Fig. 2.**
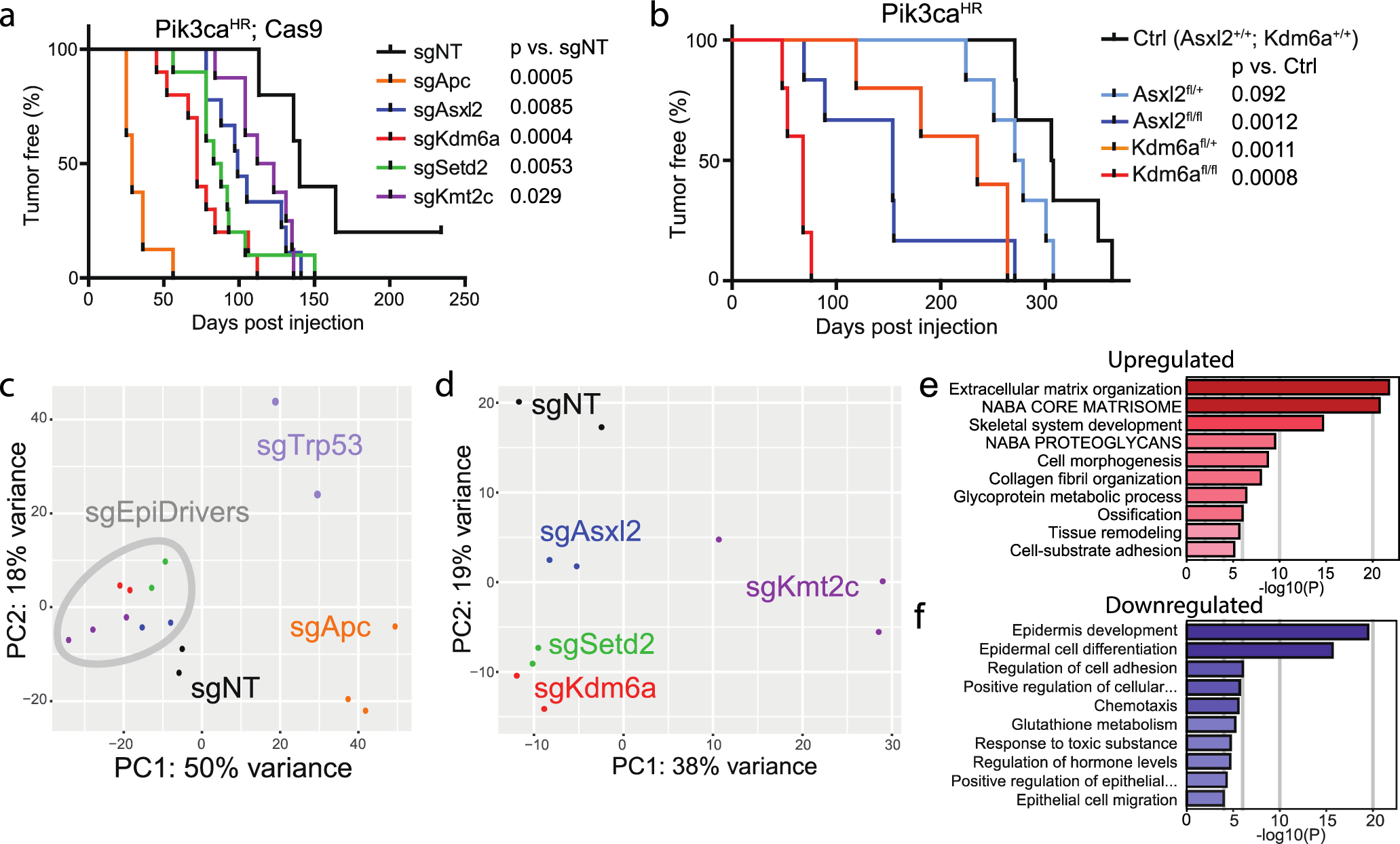
Validation and transcriptomic profiling of EpiDriver tumors. **a**, Tumor-free survival of Pik3ca^H1047R^;Cas9 mice injected with CRISPR lentivirus targeting the indicated gene or non-targeting control sgRNA (sgNT). Two independent sgRNAs/gene were used and data was combined (see Extended data Fig. 2d for single sgRNA data). **b**, Tumor-free survival of Pik3ca^H1047R^ mice with conditional knockout of Asxl2 or Kdm6a. **c**, PC plot of all profiled tumor transcriptomes. **d**, PC plot of non-targeting control and EpiDriver tumor transcriptomes. **e** and **f**, METASCAPE analysis showing enriched (**e**) and depleted (**f**) pathways in common de-regulated genes in EpiDriver-KO tumors compared to control tumors.

Histologically, mammary tumors presented as luminal-like carcinomas with either glandular, squamous, mixed squamous/glandular (adenomyoepithelioma) or spindle cell differentiation consistent with published reports of *Pik3ca^H1047R^*-induced mammary tumors (11,19). The exception was *Apc*-mutant tumors, which were exclusively adenosquamous carcinomas (**Supplementary Fig. S3A and S3B**). All tumors were estrogen receptor-positive and recapitulated gland morphology with cells marked by basal keratin 14 (K14) or luminal keratin 8 (K8). Interestingly, p53-mutant tumors showed an increased proportion of K14/K8 double positive cells, which was also seen in micro-clusters of cells invading the stroma of EpiDriver mutant tumors (**Supplementary Fig. S3C-G**).

Lastly, we generated conditional *Kdm6a^fl/fl^*;*Pik3ca^H1047R^* and *Asxl2^fl/fl^*;*Pik3ca*^H1047R^ mice and transduced the mammary epithelium with lentiviral Cre, which not only confirmed our CRISPR/Cas9 results, but also revealed that females with *Kdm6a*^fl/+^ tumors presented with significantly shorter tumor-free survival (**Fig. 2B**). *KDM6A* is located on the X-chromosome, but escapes X-inactivation and its expression reflects gene copy number (20,21). Interestingly, heterozygous *Kdm6a*^fl/+^ tumor cells still expressed Kdm6a (**Supplementary Fig. S2L and S2M**), ruling out loss-of-heterozygosity and indicating that *Kdm6a* functions as haploinsufficient tumor suppressor.

### EpiDrivers Regulate Genes involved in EMT, inflammatory pathways and differentiation

To identify pathways affected by EpiDriver loss, we obtained transcriptomes of FACS-isolated *Asxl2, Kdm6a*, *Kmt2c, Setd2, Trp53* and *Apc* knockout tumor cells. Compared to control sgNT cells, knockout tumor cells showed a wide range of differentially expressed genes (450-1800 genes; FDR <0.05, fold-change >2, **Supplemental Table S3**). Pearson’s rank correlation and principal component (PC) analysis revealed high correlation between tumors transduced with sgRNAs targeting the same gene (R^2^ >0.9) (**Fig. 2C**). Variance along PC1 and PC2 were driven by *Apc* and *Trp53* loss, respectively, suggestive of different biological impact. Consistent with the squamous histology, gene set enrichment analyses (GSEA) revealed increased transcription of genes related to keratinization in *Apc*-mutant tumors, whereas *p53*-mutant tumors showed downregulation of the p53 pathway (**Supplementary Fig. S4A and S4B**).

Unexpectedly, the PC analysis further revealed that EpiDriver tumors clustered closely with control sgNT tumors, indicating that they are transcriptionally relatively similar (**Fig. 2C**). We thus performed a PC and GSEA analysis focusing only on EpiDriver versus control sgNT *Pik3ca*^H1047R^ tumors (**Fig. 2D**). This revealed differentially upregulated pathways associated with EpiDriver inactivation including epithelial-to-mesenchymal transition (EMT), pro-inflammatory interferon-α/γ responses, downregulated cellular metabolism (oxidative phosphorylation and fatty acid metabolism) and estrogen responses (**Supplementary Fig. S4A**). We focused on 498 genes that were commonly deregulated upon loss of EpiDriver genes to elucidate potential molecular mechanisms (**Supplemental Table S4**). Pathway analysis revealed enrichment of ‘extracellular matrix organization’ and EMT and downregulation of ‘epithelial cell differentiation’ in EpiDriver tumors compared to control tumors (**Fig. 2E** **and F; Supplementary Fig. S4C and S4D**).

Intra- and cross-species comparisons revealed that control and EpiDriver knock-out *Pik3ca^H1047R^* mammary tumors also cluster together and are most similar to *Pten*-deficient mouse mammary tumors as well as human normal-like and HER2 breast cancers. This is also consistent with the fact that *PIK3CA* mutations are enriched in luminal and HER2-positive human breast cancers. In contrast, a subset of p53 knock-out tumors clustered with basal-like p53/Rb -deficient mouse mammary tumors and human basal-like breast cancer (**Supplementary Fig. S4E**). Together these data show that EpiDriver mutant tumors exhibited changes associated with EMT and block of differentiation, but generally remained histologically and transcriptionally similar to control tumors. By contrast, loss of *Apc* or *Trp53*, which showed similar reductions in tumor latency, caused dramatic transcriptional and histological changes.

### Pre-tumorigenic mammary epithelial cells display lineage plasticity and lactation mimicry

Given that the major effect of EpiDriver inactivation was accelerated *Pik3ca^H1047R^*-driven tumor initiation, we determined how their inactivation would correlate with changes in transcription and chromatin accessibility at the onset of transformation. We focused on *Kdm6a*, a core member of the COMPASS-like complex with an available conditional knock-out mouse, and performed parallel single-cell RNA sequencing (scRNA-seq) and transposase-accessible chromatin sequencing (snATAC-seq) on FACS-isolated control, *Pik3ca^H1047R^* and *Pik3ca^H1047R^;Kdm6a^fl/fl^* mammary epithelial cells two weeks after intraductal Ad-Cre injection.

We first analyzed scRNA-seq data after removing low-quality cells with low read depth (<2,500), high mitochondrial reads (>10%) and less than 1000 detected genes. This resulted in 14,070 high-quality cells composed of 6160 wild-type, 2855 *Pik3ca^H1047R^* and 5055 *Pik3ca^H1047R^;Kdm6a^KO^* cells (**Supplementary Fig. S5A**). Based on canonical markers (22), UMAP clustering revealed the three major epithelial populations corresponding to luminal progenitors (LP; *Kit*+, *Elf5*+), hormone-sensing mature luminal (HS-ML; *Prlr*+, *Pr*+, *Esr1*+) and basal cells (*Krt5/14*+) with distinct subclusters composed of the three genotypes (**Fig. 3A and B**).

**Fig. 3.**
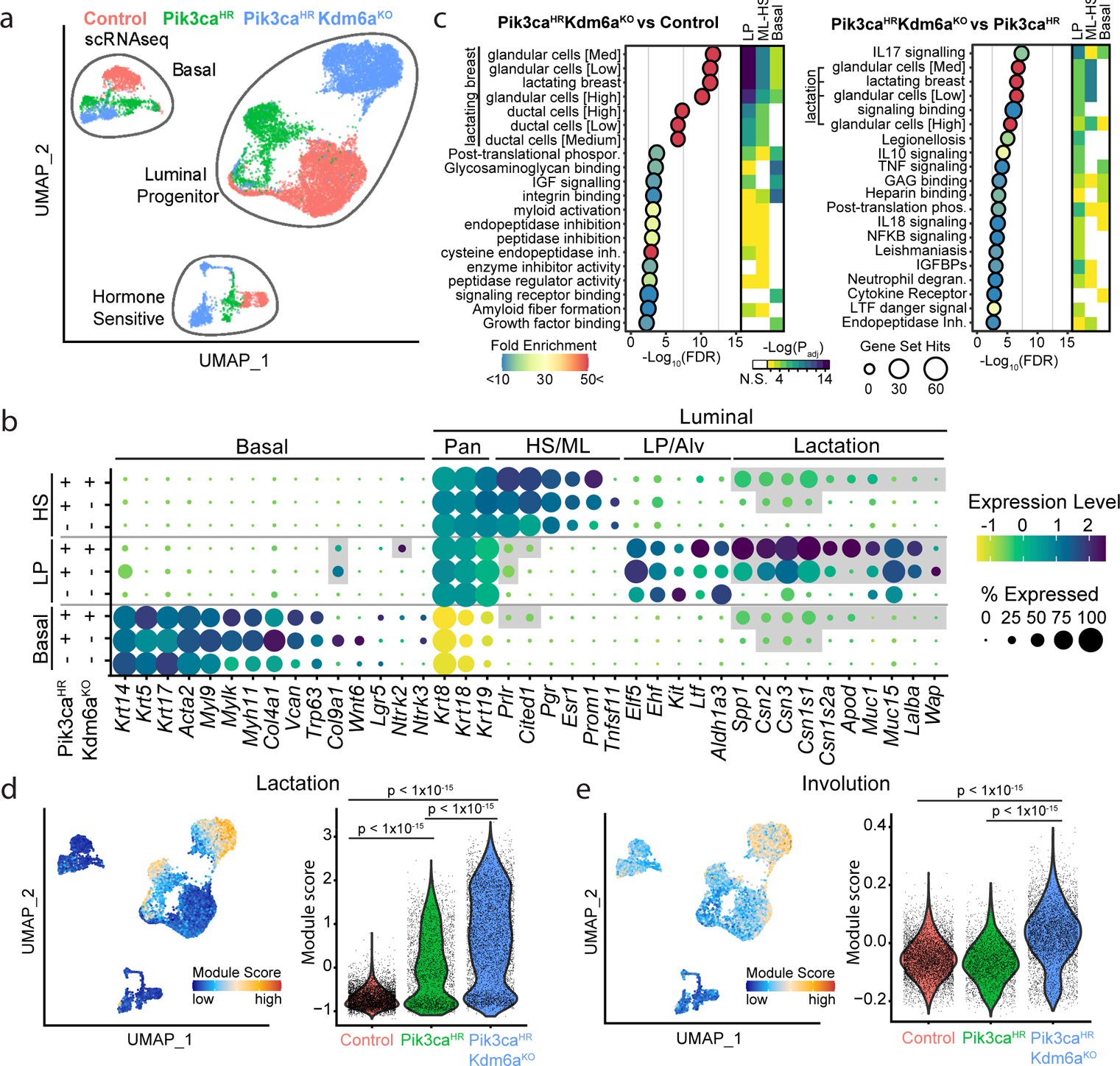
Single-cell transcriptional profiling reveals lactation mimicry. **a**, UMAP plot showing mammary epithelial cells from control, Pik3ca^H1047R^ and Pik3ca^H1047R^;Kdm6a^fl/fl^ mutant mice 2 weeks after Ad-Cre injection. **b**, Dot Blot showing differentially expressed marker genes within the different epithelial lineages stratified by genotypes. **c**, Pathways differentially enriched in Pik3ca^H1047R^;Kdm6a^fl/fl^ versus control and Pik3ca^H1047R^;Kdm6a^fl/fl^ versus Pik3ca^H1047R^ mammary epithelial LP cells identified using g:Profiler (p <0.05 with Benjamini-Hochberg FDR correction, > 10-fold enrichment). The top 20 enriched pathways are shown. Heat-map depicts how these pathways are altered in the major 3 epithelial lineages. **d** and **e**, UMAP and violin blots showing lactation **(d)** and involution **(e)** signatures.

We performed functional enrichment analysis to reveal the molecular pathways dysregulated upon activation of *Pik3ca^H1047R^* and inactivation of *Kdm6a* within each epithelial lineage. Although all our studies were performed in virgin mice, our analysis surprisingly revealed ‘lactation’ as the most differentially regulated pathway in *Pik3ca^H1047R^;Kdm6a^KO^* versus control mice, and was also highly upregulated in *Pik3ca^H1047R^;Kdm6a^KO^*versus *Pik3ca^H1047R^* mice (**Fig. 3C**). This signature was driven by genes that are normally only expressed upon differentiation of LPs into secretory alveolar cells in a hormone-dependent manner during gestation/lactation, and included caseins (*Csn2*, *Csn3*, *Csn1s1*, *Csn1s2a*), milk mucins (*Muc1/*15), lactose synthase (*Lalba*), apolipoprotein D (*Apod*) and milk proteins osteopontin (*Spp1*)*, Glycam1* and *Wap* (**Fig. 3B**). The upregulation of this lactation signature was apparent upon induction of *Pik3ca^H1047R^* but was much more prevalent upon deletion of *Kdm6a* and observed in the absence of gestation/parity-induced hormones in LP cells as well as in some basal and HS-ML *Pik3ca^H1047R^;Kdm6a^KO^* cells (**Fig. 3C and D; Supplementary Fig. S5B and S5C**).

In addition, *Pik3ca^H1047R^;Kdm6a^KO^* cells showed upregulation of genes associated with involution, EMT and hypoxia (**Fig. 3E; Supplementary Fig. S6A**). *Pik3ca^H1047R^* and *Pik3ca^H1047R^;Kdm6a^KO^* mice also exhibited higher expression of HS-ML genes such as prolactin receptor (*Prlr*) or *Cited1* in HS-ML, which also showed aberrant expression in a subset of LP and/or basal cells (**Fig. 3B; Supplementary Fig. S6B**). Conversely, basal markers such as *Krt14*, *Col9a1*, *Lgr5* and *Nrtk2* were aberrantly expressed in *Pik3ca^H1047R^* and/or *Pik3ca^H1047R^;Kdm6a^KO^* LP cells (**Supplementary Fig. S6C**). Overall, our data reveal a rapid reprogramming of the epithelial transcriptional landscape and loss of lineage integrity upon oncogenic PI3K signaling, which was exacerbated by loss of *Kdm6a*.

### snATAC-seq data confirms epigenetic re-programming and lactation mimicry

In line with the scRNAseq data and our previous data (23), unsupervised UMAP-clustering of the snATAC-seq data showed that chromatin accessibility clearly separated the three major mammary epithelial lineages (**Fig. 4A**). While wild-type, *Pik3ca^H1047R^* and *Pik3ca^H1047R^;Kdm6a^KO^* cells were intermingled in the HS-ML cluster, indicating that they are indistinguishable with regards to accessible chromatin, they formed distinct sub-clusters in the LP and to a lesser degree in the basal cluster (**Fig. 4A**).

**Fig. 4.**
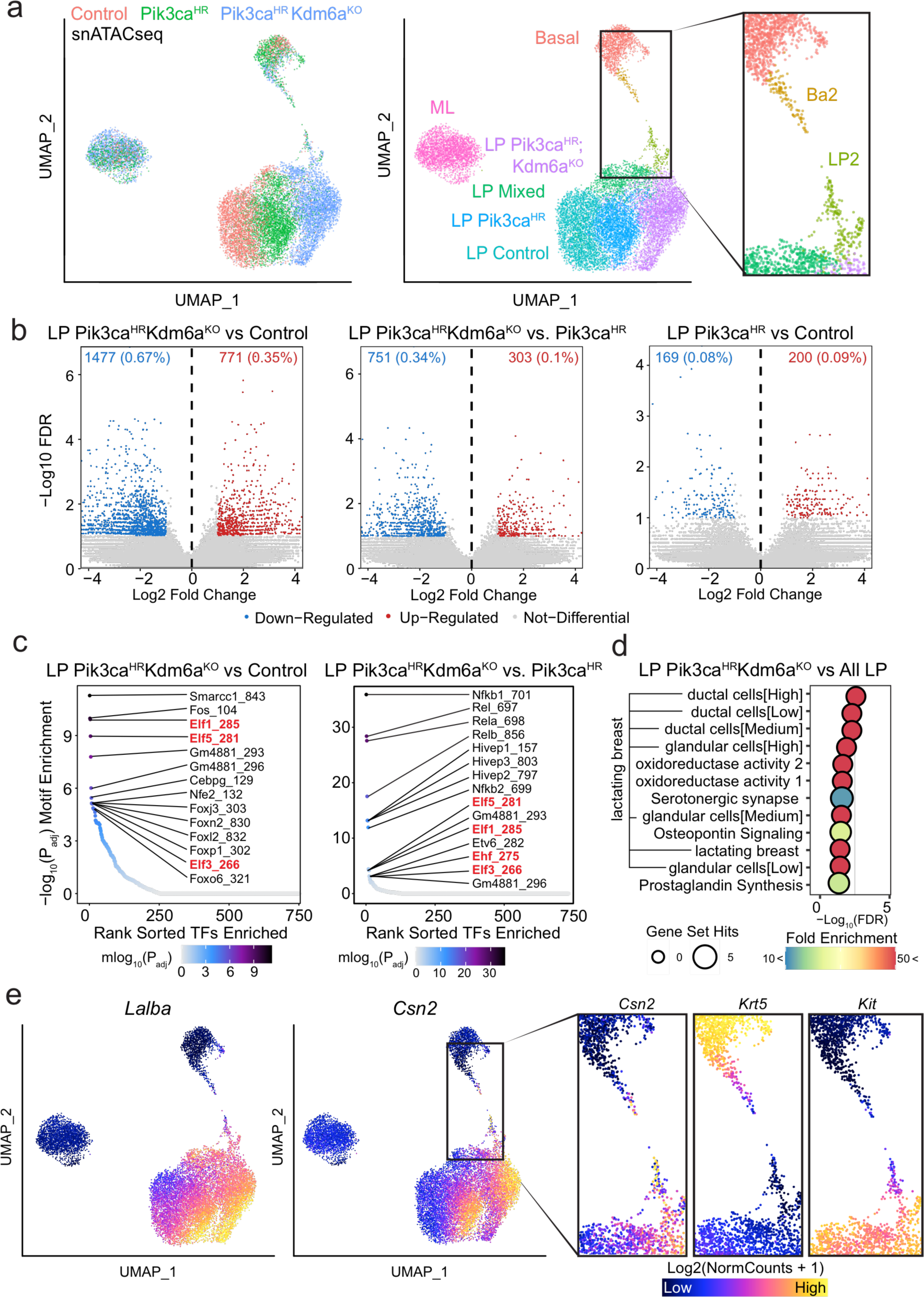
Single-cell ATACseq reveals lactation mimicry and bridge-like clusters. **a**, Unsupervised UMAP plot of snATACseq profile colored by genotype (left) and identified clusters (middle). Inlet (right) shows BA2 and LP2 clusters. **b,** Volcano plots showing differentially accessible chromatin peaks between Pik3ca^H1047R^;Kdm6a^fl/fl^ and wild-type control or between Pik3ca^H1047R^;Kdm6a^fl/fl^ and Pik3ca^H1047R^ or between Pik3ca^H1047R^ and wild-type control LP cells. **c,** Enrichment Log2(NormCounts + 1) of transcription factor binding sites in differentially accessible chromatin. **d,** Pathways differentially enriched in Log2(NormCounts + 1) Pik3ca^H1047R^;Kdm6a^fl/fl^ versus all mammary epithelial LP cells inferred from gene accessibility ArchR Gene Scores. The top 12 enriched pathways are shown as identified using g:Profiler (p <0.05 with Benjamini-Hochberg FDR correction, >10-fold enrichment). **e,** UMAP plots showing open chromatin associated with alveolar/lactation-associated genes *Lalba* and *Csn2*. Inlet (right) shows open chromatin associated with alveolar/lactation gene *Csn2*, the basal marker gene *Krt5* and the LP marker gene *Kit* in BA2 and LP2 clusters.

First, we focused on differential ATAC peaks in the LP sub-clusters, which showed the most pronounced difference between genotypes in the UMAP. While there was only a modest difference between wild-type and *Pik3ca^H1047R^* LP cells, we found a pronounced difference between *Pik3ca^H1047R^*;*Kdm6a^KO^* and wild-type as well as between *Pik3ca^H1047R^*;*Kdm6a^KO^* and *Pik3ca^H1047R^* LP cells (**Fig. 4B**), showing that loss of Kdm6a has a profound effect on chromatin accessibility. In line with Kdm6a’s H3K27 demethylase function in the COMPASS-like enhancer activation complex, we found substantially more peaks with reduced vs increased accessibility (**Fig. 4B**).

We further examined the enrichment of transcription factor motifs in the differentially accessible regions. Chromatin regions with increased accessibility in the *Pik3ca^H1047R^;Kdm6a^KO^* versus the wild-type LP cells were significantly enriched for binding sites of *Smarcc1* and *Fos* followed by ETS transcription factors *Elf1/3/5*. Motifs enriched in the *Pik3ca^H1047R^;Kdm6a^KO^* versus the *Pik3ca^H1047^* LP cells corresponded to *Nfκ-b* transcription factors *NFκ-B1/2*, *Rel* and *Rela/b* followed again by the core LP transcription factors such as *Elf1/3/5* and *Ehf* (**Fig. 4C**). Similar TF enrichment profiles were also seen in genome-wide TF activity inference using chromVAR (24) **(Supplementary Fig. S7A and S7B**). Consistent the known function of Elf5 and Ehf in driving alveolar differentiation (22,25) and in line with the scRNA-seq data, GSEA of genes associated with increased accessibility again revealed ‘lactation’ as the most significant gene sets upregulated in *Pik3ca^H1047R^;Kdm6a^KO^* LP cells with increased accessibility of multiple alveolar/milk biogenesis genes such as *Csn2/1s1/1s2a, Lalba, Apod, Spp1, Lipa, Lif* (**Fig. 4D and E; Supplementary Fig. S8A**).

In addition, UMAP further revealed a basal-like ‘Ba2’ and a luminal-like ‘LP2’ subcluster enriched in *Pik3ca*^H1047R^ and *Kdm6a^KO^;Pik3ca^H1047R^* cells that appear to bridge the basal and LP populations (**Fig. 4A**). GSEA of genes associated with increased accessibility in Ba2/LP2 cells revealed gene sets associated with ‘chromatin silencing’ (*Sox12, Sin3a, Adat2*, histones *H3c1, H4c1* etc.), ‘FOXO-mediated transcription of oxidative stress’ (*Sox8, Ltbp4, Egr1*) and ‘epithelial tube morphogenesis’ (*Dvl2, Fzd2, Dlg4*, *Prickle4*) (**Supplementary Fig. S8A and S8B**). In addition, the biological KEGG pathway ‘breast cancer’ was upregulated in the Ba2 versus the basal cluster with prominent WNT signaling genes such as *Wnt10a, Wnt6, Fzd2, Dvl2, Prickle4, Csnk1g2* and *Dlg4* or NOTCH signaling such as *Dll1* and *Jag2* (**Supplementary Fig. S8A and S8C**). In line with this notion, chromVAR analysis showed enrichment of binding sites for transcription factors associated with WNT (*Lef1, Tcf7, Tcf7l1, Tcf7l2*) and NOTCH signaling (*Hes1, Hey1/2, Heyl*) in Ba2 cells (**Supplementary Fig. S7A**). Consistently, we observed upregulation of WNT and NOTCH signaling signatures in *Pik3ca^H1047R^* and *Pik3ca^H1047R^;Kdm6a^KO^* basal cells in the parallel scRNA-seq dataset (**Supplementary Fig. S8D and S8E**). Of note, *Apc* was the major hit in the *in vivo* CRISPR screen (**Fig. 1F**), suggesting that elevated WNT signaling is oncogenic in our *Pik3ca^H1047R^* model. In addition, WNT and NOTCH signaling are not only known drivers of breast cancer but also play critical roles in mammary lineage determination (26-28).

Overall, we found that Ba2 cells have reduced chromatin accessibility at basal markers such as *Krt5/14*, *Trp63*, *Acta2* (=smooth muscle actin) or vimentin and increased accessibility of the alveolar genes *Csn2*, whereas LP2 cells have reduced chromatin accessibility at LP markers such as *Elf5, Ehf*, and *Kit* (**Fig. 4E; Supplementary Fig. S9A-C and S10A-C**), consistent with loss of lineage identity observed in the scRNAseq data. Together, our scRNAseq and snATACseq data suggest that *Pik3ca^H1047R^;Kdm6a^KO^* mammary epithelial cells gain lineage plasticity and prior to tumorigenesis reprogram towards the alveolar fate reminiscent of lactation.

All scRNAseq data are accessible in this <u>interactive Shiny application</u>.

### The COMPASS-like complex inhibits a tumorigenic basal-to-luminal cell lineage conversion

We next determined whether both luminal and basal cells are susceptible to lineage plasticity and contribute to tumor formation. We used lineage tracing with a basal-specific adenoviral Ad-K5-Cre and luminal-specific Ad-K8-Cre viruses (29) to address this question (**Supplementary Fig. S11A-S11E)**. As previously shown (30)(31), expression of oncogenic *Pik3ca^H1047R^* can lead to lineage plasticity and convert basal and luminal unipotent progenitors into multipotent cells. In line with these reports, induction of *Pik3ca^H1047R^* in basal cells resulted in a gradual lineage conversion to luminal-like cells, which was dramatically accelerated by *Kdm6a* or *Asxl2* deletion (**Fig. 5A-C; Supplementary Fig. S11D**). In line with the haploinsufficiency, heterozygous loss of *Kdm6a* also significantly accelerated linage conversion (**Supplementary Fig. S11F**). In contrast to prior studies, we did not observe accelerated lineage conversion from luminal-to-basal cells (**Supplementary Fig. S11G**).

**Fig. 5.**
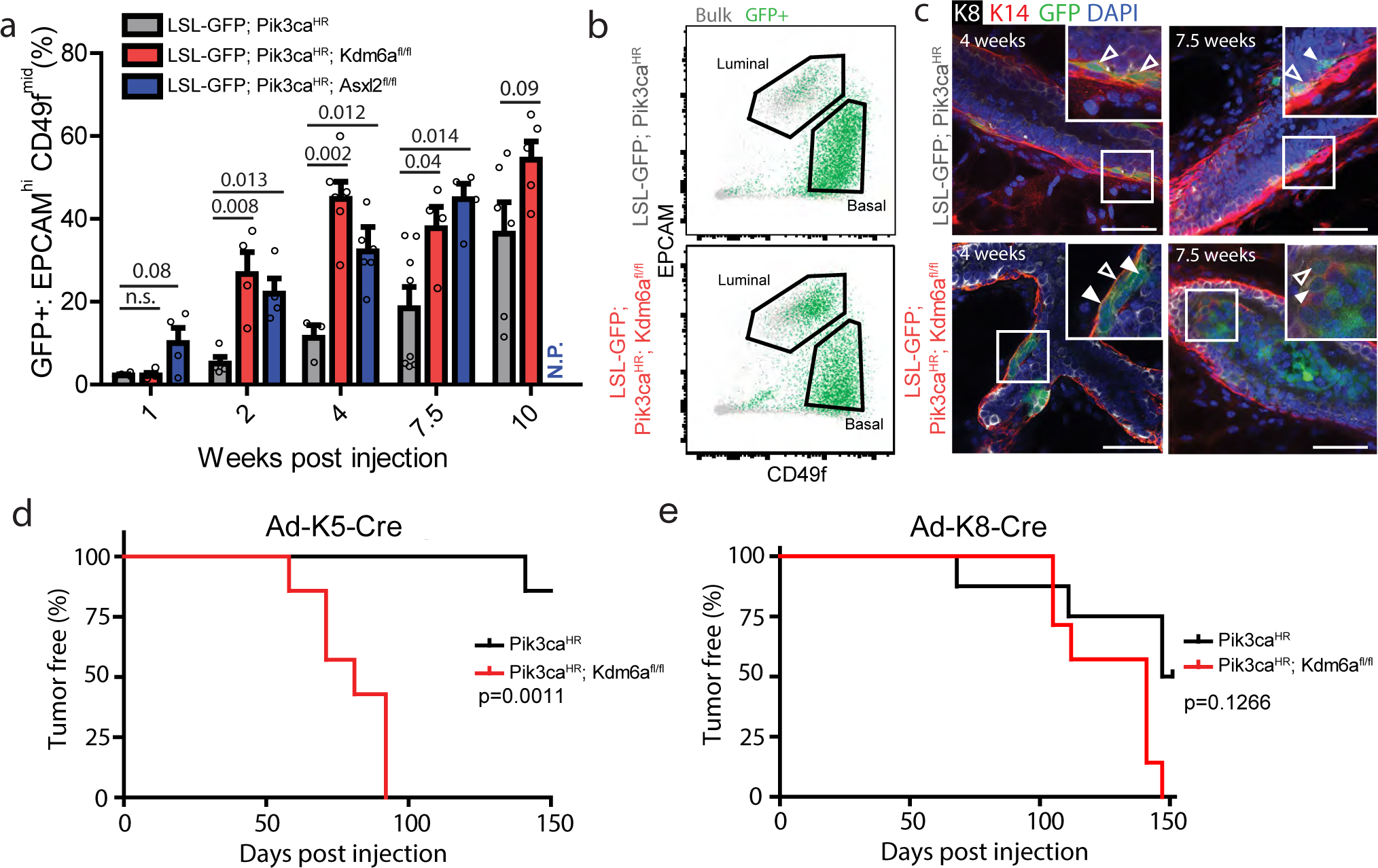
Loss of EpiDrivers induces multipotency. **a**, Percent of GFP+ EPCAM^high^ CD49f^mid^ luminal cells at different time points after Ad-K5-Cre injection into mammary epithelium of mice with the indicated genotype. **b**, Representative FACS plot at 4 weeks post injection with Ad-K5-Cre. **c**, Whole-mount image of mammary glands 4 weeks and 7.5 weeks post Ad-K5-Cre injection showing K14+/K8- (empty arrows) as well as K14+/K8+ double-positive and K14-/K8+ GFP+ lineage-traced cells (filled arrows). Scale bar = 50 µm. **d** and **e**, Tumor-free survival of Pik3ca^H1047R^;Kdm6a^fl/fl^ versus Pik3ca^H1047R^ after intraductal injection of Ad-K5-Cre (**d**) and Ad-K8-Cre (**e**).

Next, we used the K5- and K8-Cre drivers to determine if the cell-of-origin affects the latency and phenotype of tumors arising in *Pik3ca^H1047R^;Kdm6a^fl/fl^* mice. Loss of *Kdm6a* in the basal compartment significantly accelerated tumor formation, whereas luminal cell-derived *Pik3ca^H1047R^;Kdm6a^KO^* tumors arose with similar latency as *Pik3ca^H1047R^* tumors (**Fig. 5D and E**). Transcriptome analysis revealed that basal-cell derived tumors were indistinguishable from those derived upon sgRNA-mediated mutation of *Kdm6a* (**Supplementary Fig. S4C**). Together, these results indicate that loss of the COMPASS-like complex in *Pik3ca^H1047R^* basal cells accelerates their reprogramming into tumor-initiating cells that drive luminal-like breast cancer.

To further characterize the basal-to-luminal-like cell transition, we performed scRNA-seq on control, *Pik3ca^H1047R^* or *Pik3ca^H1047R^;Kdm6a^KO^* mammary epithelial cells after two weeks of Ad-K5-Cre lineage-tracing (**Fig. 6A-D; Supplementary Fig. S12A and S12B**). Consistent with the results above, LP-like cells that lost basal markers and gained LP (*Kit, Elf5, Cd14 or Aldh1a3*) and alveolar markers (*Ehf*, *Cns3, Apod* or *Wfdc18)* emerged from *Pik3ca^H1047^* and *Kdm6a^KO^;Pik3ca^H1047R^* basal cells. We even observed rare *Pik3ca^H1047^* and *Pik3ca^H1047R^;Kdm6a^KO^* cells expressing milk genes such as *Wap* and *Olah* or HS-ML markers such as *Prlr* (**Fig. 6E-G; Supplementary Fig. S12B, S12C and S13A-C**).

**Fig. 6.**
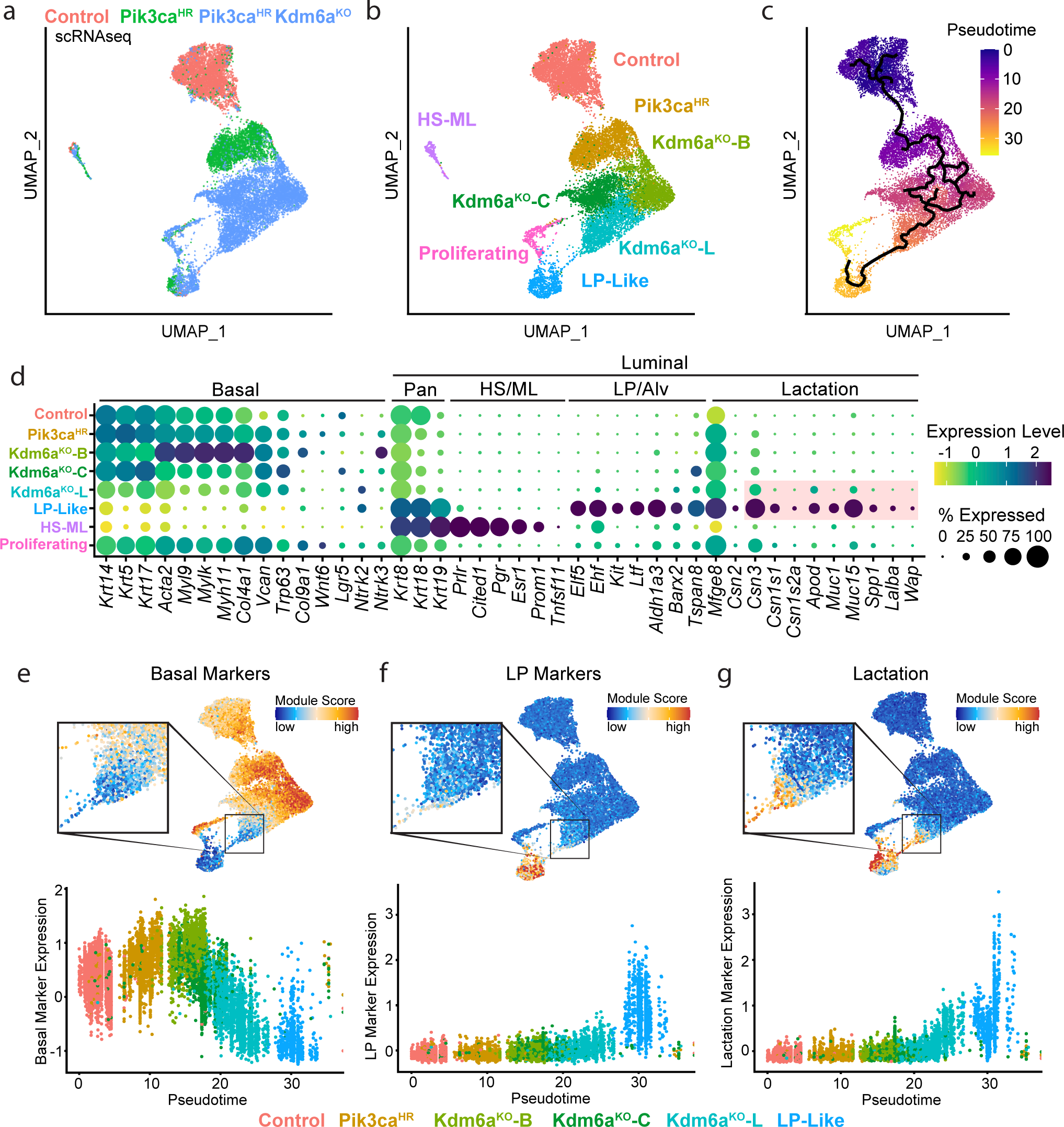
scRNAseq reveals basal-to-alveolar transdifferentiating at the onset of breast cancer initiation. **a-c**, UMAP plots showing Ad-K5-Cre lineage-traced basal mammary epithelial cells from control, Pik3ca^H1047R^ and Pik3ca^H1047R^;Kdm6a^fl/fl^ mutant mice 2 weeks post-injection colored by genotype (**a**), clusters (**b**) and trajectories inferred by Monocle3 (**c**). **d**, Dot plot showing differentially expressed marker genes within the different epithelial clusters. **e-g**, UMAP and pseudo-time trajectory plots showing basal (**e**), luminal progenitor (**f**) and alveolar/lactation (**g**) marker signatures.

In addition, *Pik3ca^H1047R^;Kdm6a^KO^* basal cells were more heterogenous than wild-type or *Pik3ca^H1047^* cells and comprised three unique subclusters: Kdm6a^KO^-L, adjacent to the <u>L</u>P-like population, a <u>c</u>entral cluster (Kdm6a^KO^-C), and a cluster enriched in <u>b</u>asal markers (Kdm6a^KO^-B; *Myh11, Myl9, Acta2, Igfbp2*) further underscoring the notion of increased phenotypic plasticity upon loss of *Kdm6a*. Importantly, Kdm6a^KO^-L showed a gradual downregulation of basal markers with concomitant upregulation of alveolar/lactation markers such as *Csn2/3, Apod*, *Muc1/15* or *Wfdc18* (**Fig. 6E-G; Supplementary Fig. S12B, S13A-C and S14A-C)**. Kdm6a^KO^-L was also marked by expression of the EMT master regulator *Zeb1* and *Zeb2*, *PTN*, *Sox10*, *Tgfb2* and the latent TGFB binding protein *Ltbp1* as well as Suppressor Of Cytokine Signaling *Socs2*, and Neurotrophic Receptor Tyrosine Kinase 2 *Ntrk2* (**Supplementary Fig. S14D**). Of note, *Ntrk2* was previously identified as a basal-to-luminal multipotency breast cancer gene (30) and together with *Ptn* are known breast cancer promoting genes (32). Interestingly, this Kdm6a^KO^-L cluster did not express classic luminal progenitor markers (*Kit, Elf5, Aldh1a3, Cd14, Lif*) (**Fig. 6F; Supplementary Fig. S12C**). This observation combined with trajectory analysis (**Fig. 6C**) suggests that *Kdm6a^KO^;Pik3ca^H1047R^* basal cells start to gradually activate an aberrant alveolar-like program before acquiring LP characteristics.

Integrating the Ad-Cre and the Ad-K5-Cre scRNAseq datasets revealed that the luminal-like Ad-K5Cre-lineage-traced *Pik3ca^H1047^* and *Pik3ca^H1047R^;Kdm6a^KO^* cells clustered with LP cells from the first Ad-Cre scRNAseq experiment, further supporting the notion of a basal-to-luminal reprogramming. In addition, luminal-like Ad-K5Cre-lineage-traced *Pik3ca^H1047^* and *Pik3ca^H1047R^;Kdm6a^KO^* with a high lactation and involution signature clustered with *Pik3ca^H1047^* and *Pik3ca^H1047R^;Kdm6a^KO^* LP cells, while those without a lactation/involution signature clustered with wt LP cells, indicating functional heterogeneity (**Supplementary Fig. S15A-D)**.

Cells in the proliferating cluster consisted mainly of *Pik3ca^H1047R^;Kdm6a^KO^* with either basal or luminal characteristics (**Fig. 6A, B and E-F**). This cluster also showed marked elevation of RB1/E2F target genes such as *Brca1/2, Mcm2/6, Pole, Chek1, Chaf1b, Cdca8, Exo1* or *Ect2* (**Supplementary Fig. S16**), reminiscent of the RB1 inactivation and E2F activation during pregnancy-induced hyperproliferation in the mammary gland (33), further supporting a role of these cells and the aberrant alveolar program during tumor initiation.

### EpiDrivers function as tumor suppressors in human breast cancer

To extend our findings from mouse to human cancers, we assessed the function of the EpiDrivers in human MCF10A mammary epithelial cells that harbor a *PIK3CA^H1047R^* knock-in mutation(34,35). Using CRISPR/Cas9, we generated *ASXL2*, *KDM6A, KMT2C*, *SETD2, PTEN* and *TP53* knock-out lines as well as control sgNT cells (**Supplementary Fig. S17A and S17B**). Like parental MCF10A cells, *PIK3C^AH1047^*^R^ cells formed polarized and hollow, albeit a little larger, acini in Matrigel culture^22^. In contrast, *ASXL2*-, *KDM6A*-, *KMT2C*-, *PTEN-*, and *TP53-*mutant spheres showed a transformed phenotype of with large branching protrusions (**Fig. 7A** and **B**). When grafted orthotopically into the fat pads of NSG mice, s*gKDM6A, sgSETD2, sgTP53* and sg*PTEN* cells formed tumors while control sgNT cells did not (**Fig. 7A; Supplementary Fig. S17C**). Although *sgASXL2* and sg*KMT2C* cells exhibited a transformed phenotype in 3D cultures, they did not efficiently give rise to xenograft tumors in mice. Together, these data show that *KMT2C, KDM6A, ASXL2* and *SETD2* suppress transformation of human MCF10A mammary epithelial cells.

**Fig. 7.**
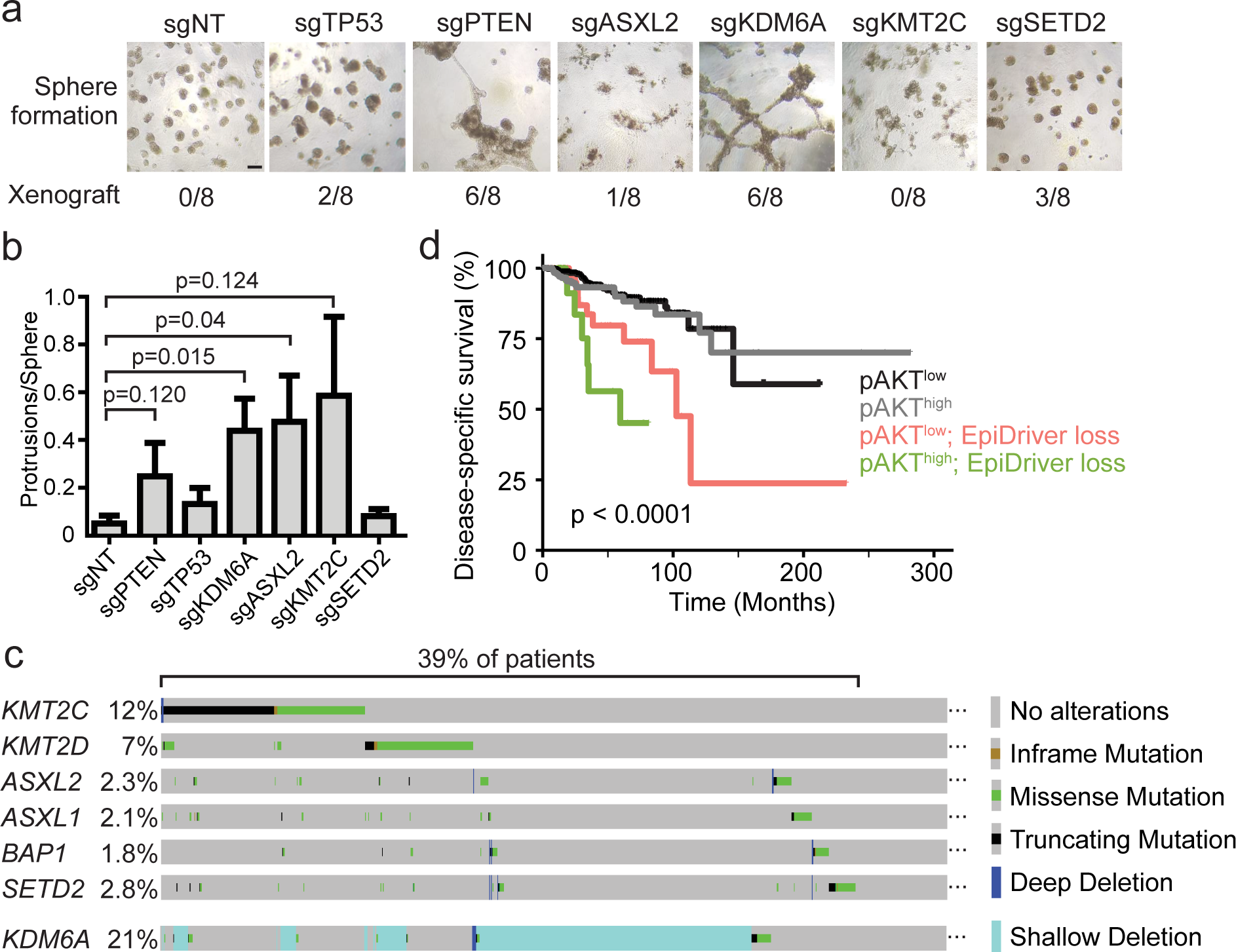
EpiDrivers function as Tumor Suppressors in Humans. **a**, Representative images of spheres formed by MCF10A-PIK3CA^H1047R^ cells with CRISPR/Cas9 knockout of the indicated genes (top) and their xenograft efficiency in NSG mice (bottom). **b**, Quantification of number of sphere protrusions of MCF10A-PIK3CA^H1047R^ cells with indicated EpiDriver loss. **c**, Prevalence of alterations in EpiDrivers in human breast tumors. Shallow deletion only displayed for KDM6A. **d**, Disease-specific survival (DSS) of breast cancer patients in the TCGA cohort stratified by phospho-Ser473 AKT (pAKT) and EpiDriver mutations. The long-rank p value is shown.

Next, we analyzed 2443 METABRIC breast cancer cases (10,12). *ASXL2, BAP1, KDM6A, KMT2C*, *KMT2D,* and *SETD2* are mutated in 1-12% of breast tumors, as expected for long-tail genes (**Fig. 7C; Supplementary Fig. S18A**) (10,12). The haploinsufficiency of *Kdm6a* in mouse mammary tumorigenesis prompted us to also analyze copy number alterations. Interestingly, an additional 19% of patients exhibited shallow deletion indicative of heterozygous *KDM6A* loss (**Fig. 7C**), which coincided with significantly reduced *KDM6A* expression (**Supplementary Fig. S18B**). In addition, EpiDriver alterations showed a trend towards mutual exclusivity but displayed a considerable overlap with *PIK3CA* mutations (**Supplementary Fig. S18A**). Survival analysis of the TCGA breast cancer cohort showed that tumors with an EpiDiver mutations and high PI3K signaling defined by high phospho-Ser473-AKT (36) or an high transcriptional PI3K signature (37) stratified patients with the poorest survival across subtypes (**Fig. 7D**) as well as within Luminal A and B breast cancer (**Supplementary Fig. S18C-S18E**). About 39% of patients harbor alterations in the KMT2C/D-COMPASS-like or BAP1/ASXL complexes (**Fig. 7C**), highlighting the importance of this tumor suppressive network.

## Discussion

Large international efforts such as <u>TCGA</u> and <u>ICGC</u> have set out to profile the mutational landscape of many cancers with the goal of cataloguing the genes responsible for tumor initiation and progression. The idea was to identify those genes that are mutated more frequently than expected by random chance and the expectation was that increasing sample size will boost the power to mathematically infer real driver mutations (=sensitivity), while weeding out background of random mutations (=specificity). However, recent studies show that the more we sequence, the more seemingly statistical genes we identify giving rise to the ‘long tail’ of cancer associated genes. This suggest that despite sequencing hundreds to thousands of tumors of a given cancer type, we have not yet reach saturation. In addition, many of the lower frequency genes seem implausible such as olfactory receptors, suggesting that the mutational processes inherent to carcinogenesis (noise) overshadow true driver events (signal), giving rise to a high false-positive rate. Further increasing sample size to characterize the observed mutational heterogeneity will only marginally improve signal-to-noise.

Here, we devised and deployed an *in vivo* CRISPR/Cas9-screening methodology, which allowed us to identify *bone-fide* cancer driver mutation in the long tail of breast cancer. Our *in vivo* CRISPR/Cas9-screen identified several tumor suppressor genes in the long-tail of breast cancer-associated mutations with our top hits converging on epigenetic regulation. Individually, epigenetic regulators are not mutated frequently, but as a group, they are among the most frequently mutated targets in cancer (38-42), indicating that a ‘deregulated epigenome’ can accelerate tumor development. In particular, we identified several components and auxiliary factors of the COMPASS-like histone methyltransferase complex as potent tumor suppressors and showed that *Kdm6a* might function in a haploinsufficient manner. Up to 39% of breast cancer patients harbor mutations in the COMPASS-like pathway, highlighting the importance of elucidating the mechanisms by which COMPASS inactivation contributes to breast cancer progression.

Components of the COMPASS-like complex were recently implicated as tumor suppressors in leukemia (43), medulloblastoma (44), pancreatic (21) and non-small-cell lung cancer (45) and their loss was associated with substantial enhancer reprogramming and aberrant transcription. We were surprised to find that EpiDriver inactivation did not substantially affect histology or transcriptional profiles of breast tumors. However, it did significantly accelerate tumor initiation, which was coupled with rapid acquisition of phenotypic plasticity. Plasticity plays a central role in development and during tissue regeneration and wound healing (23,46,47). More recently, phenotypic plasticity has also been recognized as a driving forces behind tumor initiation and progression (48-50). For example, elegant lineage-tracing and single cell-profiling experiments have shown that oncogenic signaling can reactivate multipotency within the two epithelial lineages of the mammary gland (30,31,48). Cells that acquire plasticity are thought to gain stem cell features through a process of dedifferentiation (47,51). However, in the system studied here, we did not observe acquisition of fetal mammary stem cell-like transcriptomes as observed in basal-like tumors studies (23,48). Rather, we observed an aberrant differentiation program associated with alveologenesis and lactation induced upon Pik3ca activation and exacerbated by EpiDriver loss. This was most noteworthy in basal cells, which are known to be functionally plastic (52-54). A similar aberrant alveolar differentiation program was recently described in breast cancer models driven by luminal loss of Brca1 and p53 (22) and upon luminal overexpression of Elf5 and PYMT (55,56). Together, our data indicate that there are different avenues towards transformation and that the innate but poised program coordinating the proliferative burst during gestation/lactation can be being highjacked for rapid expansion at the onset of oncogenic transformation – a phenomenon we term lactation mimicry. This lactation mimicry is dramatically exacerbated/accelerated by loss of epigenetic control governed by the COMPASS-like complex and associated BAP1/ASXL1/2 complex and happens not only in the luminal cells, but – given the right combinations of mutations – also in the basal cells. It will be interesting to assess whether other cancers also coerce regenerative or tissue remodeling process inherent to that tissue during early transformation.

Another interesting aspect of our study is the potential cell of origin underlying different subtypes of breast cancer. Gene expression studies indicated that mature luminal cells give rise to luminal A/B and HER2 subtypes, while luminal progenitors transform to the basal-like cancers and basal cells give rise to the claudin-low subtype (57-59). Mouse lineage-tracing studies have supported these observations and have shown that certain mutations in specific lineages can indeed give rise to mouse mammary tumors with features similar to different human breast cancer subtypes (30,31,60). Our data now show that, given the right combination of oncogene and cooperating epigenetic alteration, basal cells can also be the cell of origin of luminal tumors. This supports the idea that the ultimate epigenomic, transcriptomic, and histopathologic characteristics of a tumor depend on the target cell for the initial mutation, the type of mutations, and the collaborating alterations. Clearly, loss of epigenetic regulation needs to be considered as a significant contributor to the loss of lineage integrity that underlie intra-tumoral heterogeneity.

## Data Availability

All RNA-seq, scRANseq and snATACseq are available at NCBI Gene Expression Omnibus GEO accession GSE178424. The following secure token has been created to allow review of record GSE178424 while it remains in private status. This token allows anonymous, read-only access to GSE178424 and associated accessions while they are private. To review GEO accession GSE178424 go to: https://www.ncbi.nlm.nih.gov/geo/query/acc.cgi and enter token **mterweoyzfozfgj** into the box.

## Acknowledgements

We thank all members of our laboratories for helpful comments, with additional thanks to Y.Q. Lu, K. Schleicher, and G. Mbamalu for their insight and assistance. We thank H. Melo and D. Durocher for assistance with visualization of g:Profiler data. We also thank The Centre for Phenogenomics, Network Biology Collaborative Centre and Flow Cytometry facility at LTRI as well as the Flow Cytometry Facility at the University of Toronto. Funding: This work was supported by a Career Catalyst Research Grant to D.S. from Susan G. Komen foundation (CCR16377321), a Terry Fox Research Institute Program Projects Grant to J. W. and D.S et al. (TFRI Project #1107) and by Nicol Family Foundation. E.L is a recipient of the Ontario Graduate Scholarship and the Frank Fletcher Memorial Fund, K.N.A. is a recipient of the MBD fellowship and supported by SHSF donation (Mr. Ah Shai), S.K.L. is a Canadian Cancer Society Fellowship recipient (BC-F-16#31919). GMW and ZM are supported by a Cancer Center Core Grant (5 P30CA014195), NIH/National Cancer Institute (R35 CA197687), and the Breast Cancer Research Foundation (BCRF). S.E.E is supported by CIHR. R.P. and M.A.P. was supported by grants from the Carlos III Institute of Health (PI18/01029; co-funded by European Regional Development Fund (ERDF), a way to build Europe), Generalitat de Catalunya (SGR 2017-449), and the CERCA Program to IDIBELL.

## Author Contributions

E.L. performed all experiments. K.N.A., S.K.L., R.T. and T.N. helped with mouse experiments and FACS analysis, A.M., J.L, H.W.J., G.B. and M.A.P performed bioinformatics analysis. D.T and J.W performed the scRNAseq and snATACseq experiment. Z.M. and G.M.W analyzed all the single cell sequencing data, K.K and S.E.E performed histological analyses. R.B, E.S.K and H.W.J helped with experimental design. D.S. coordinated the project and, together with G.M.W and E.L designed the experiments and wrote the manuscript.

## Supplementary Materials

### Methods

#### Animals

Animal husbandry, ethical handling of mice and all animal work were carried out according to guidelines approved by Canadian Council on Animal Care and under protocols approved by the Centre for Phenogenomics Animal Care Committee (18-0272H). The animals used in this study were R26-LSL-Pik3caH1047R/+ mice (11) [Gt(ROSA)26Sortm1(Pik3ca*H1047R)Egan in a clean FVBN background kindly provided by Egan S, SickKids], R26-LSL-Cas9-GFP [#026175 in C57/Bl6 background from Jackson laboratories], LSL-TdTomato [#007908 from Jackson laboratories], Asxl2fl/fl [C57BL/6N-Asxl2<tm1c(EUCOMM)Hmgu>/Tcp generated by The Canadian Mouse Respiratory] and Kdm6afl/fl [Kdm6atm1.1Kaig] mice kindly provided by Jacob Hanna. CRISPR screens in the Pik3caH1047R/+; Cas9 cohort were performed in a F1 FVBN/C57Bl6 background. Genotyping was performed by PCR using genomic DNA prepared from mouse ear punches. When total tumor mass per animal exceeded 1000mm^3^, mice were monitored bi-weekly and scored in accordance to SOP “#AH009 Cancer Endpoints and Tumour Burden Scoring Guidelines”.

#### Lentiviral constructs and library construction

sgRNAs targeting breast cancer long tail genes were obtained from Hart et al. (61) (4 sgRNAs/gene) and non-targeting sgRNAs were obtained from Sanjana et al. (62), ordered as a pooled oligo chip (CustomArray Inc., USA) and cloned into pLKO sgRNA-Cre plasmid (8) using BsmBI restriction sites. We excluded frequent and known breast cancer tumor suppressor genes such as *TP53* or *CDH1* from the breast long tail genes library. The non-targeting sgRNAs are those designed not to target the mouse genome as negative control. pLKO-mRFP and pLKO-GFP were kindly provided by Elaine Fuchs (Addgene #26001 and #25999). pLKO-mRFP-P2A-Cre was recently described (8) and used for lentiviral injections in *Pik3ca*^H1047R^;*Kdm6a*^fl/fl^ and *Asxl2*^fl/fl^ mice. Ad-Cre, Ad5-K5-Cre and Ad5-K8-Cre was purchased from the Vector Core at the University of Iowa.

#### Lentivirus production and transduction

Large-scale production and concentration of lentivirus were performed as previously described (63-67). Briefly, 293T cells (Invitrogen R700-07) were seeded on a poly-L-lysine coated 15 cm plates and transfected using PEI (polyethyleneimine) method in a non-serum media with lentiviral construct of interest along with lentiviral packaging plasmids psPAX2 and pPMD2.G (Addgene plasmid 12259 and 12260). 8 hours post-transfection media was added to the plates supplemented with 10% Fetal bovine serum and 1% Pencillin-Streptomycin antibiotic solution (w/v). 48 hours later, the viral supernatant was collected and filtered through a Stericup-HV PVDF 0.45-μm filter, and then concentrated ∼2,000-fold by ultracentrifugation in a MLS-50 rotor (Beckman Coulter). Viral titers were determined by infecting R26-LSL-tdTomato MEFs and FACS based quantification. *In vivo* viral transduction efficiency was determined by injecting decreasing amounts of a single viral aliquot of known titer, diluted to a constant volume of 8 μl per mammary gland. We collected mammary glands at 7 days post-infection and determined percent infection using FACS.

#### Intraductal injection and viral transduction

Intraductal lentiviral injection have been described. Briefly, to deliver the lentiviral sgRNA library or single sgRNAs targeting gene of interest, a non-invasive, injection method was employed, which selectively transduces mammary epithelium of female mice. Female mice were injected at >8 and <20 weeks of age, with age at injection matched between groups in all experiments. 8 ul of virus diluted in PBS and visualized with Fast-Green dye was injected into the 3^rd^ and/or 4^th^ mammary glands using pulled glass micropipettes. As previously described (63,65,67), we calculated coverage based on the following parameters: mammary epithelium consist of ∼3.5x10^5^ cells; transduction of ∼15% results in a minimal double infection rate (∼1/10 infected cells); at 15% infectivity every gland has 50.000 infected cells and 200.000 cells in four glands of a single mouse; To ensure that at least 4000 individual cells were transduced with a given sgRNA, a pool of 860 sgRNAs requires 3.5x10^6^ cells or ∼17 animals. To verify the sgRNA abundance and representation in the control and breast long-tail genes libraries, MEFs were transduced with library virus and collected 48h post transfection. For hit validation, lentivirus was injected at 1x10^7^ pfu/ml. Ad5-K5-Cre virus was injected at 8x10^8^ pfu/ml and Ad-K8-Cre virus was injected at 3.5x10^10^ pfu/ml, which infected ∼2-10% of basal or luminal cells respectively.

#### Deep Sequencing: sample preparation, pre-amplification and sequence processing

Genomic DNA from epithelial and tumor cells were isolated with the DNeasy Blood & Tissue Kit (Qiagen). 5μg genomic DNA of each tumor was used as template in a pre-amplification reaction using unique barcoded primer combination for each tumor with 20 cycles and Q5 High-Fidelity DNA Polymerase (NEB). The following primers were used: FW:5’AATGATACGGCGACCACCGAGATCTACAC**<u>TATAGCCT</u>**ACACTCTTTCCCTACACGACGCTCTTCCGATCTtgtggaaaggac gaaaCACCG-3’ RV:5’CAAGCAGAAGACGGCATACGAGAT**<u>CGAGTAAT</u>**GTGACTGGAGTTCAGACGTGTGCTCTTCCGATCTATTTTAACTTG CTATTTCTAGCTCTAAAAC-3’

The underlined bases indicate the Illumina (D501-510 and D701-712) barcode location that were used for multiplexing. PCR products were run on a 2% agarose gel, and a clean ∼200bp band was isolated using Zymo Gel DNA Recovery Kit as per manufacturer instructions (Zymoresearch Inc.). Final samples were quantitated then sent for Illumina Next-seq sequencing (1 million reads per tumor) to the sequencing facility at Lunenfeld-Tanenbaum Research Institute (LTRI). Sequenced reads were aligned to sgRNA library using Bowtie version 1.2.2 with options –v 2 and –m 1. sgRNA counts were obtained using MAGeCK count command.

#### Analysis of genome editing efficiency

LSL-Cas9-GFP MEFs were cultured and infected with lentivirus carrying Cre and corresponding sgRNAs. Cells were live sorted for GFP expression and expanded further to extract genomic DNA using DNeasy Blood & Tissue Kit (Qiagen). Genomic DNA from tumors from the mice injected with single sgRNAs were also isolated using the same kit. PCR was performed flanking the regions of sgRNA on genomic DNA from both WT MEFs and cells infected with respective virus or tumors and sent for Sanger sequencing. Sequencing files along with chromatograms were uploaded to https://www.deskgen.com/landing/tide.html and genome editing efficiency was estimated. sgRNAs and TIDE primers are listed in Supplementary Table S5.

#### Antibodies

The following primary antibodies were used in this study: rabbit anti-APC (1:200, Santa Cruz sc-896), rabbit anti-Kdm6a (1:1000, CST D3Q1I), rabbit anti-Asxl2 (1:500, EMD Millipore), mouse anti-TP53 (1:1000, CST 1C12), mouse anti-Pten (1:1000 CST 26H9), goat anti-Setd2 (1:500 Millipore-Sigma SAB2501940), rabbit anti-Mll3 (1:500 CST D1S1V), mouse anti-GAPDH (1:2500 Santa Cruz sc-32233), rabbit anti-histone H3 (1:1000 CST 4499), rabbit anti-Keratin14 (PRB-155P, 1:200 for whole mount, 1:700 for sections), rat anti-Keratin8 (1:50, TROMA-1), APC conjugated antiCD45, (1:500 rat monoclonal Clone 30 F11), APC conjugated antiCD31 (1:250 rat monoclonal Clone MEC133, Biolegend), APC AntiMouse Ter119 (1:250 Biolegend), PECy7 anti human/mouse CD49f (1:50 clone GoH3, Biolegend), APCVio770 mouse anti-CD326 EpCAM (1:50 Miltenyi).

#### Cell culture

Primary mouse tumor cells were cultured in DMEM/F12 (1:1) supplemented with MEGS supplement, FBS and Pen Strep. MCF10A cells were cultured as previously described (68). Cells were cultured in monolayer for growth and transfection with lentiviral CRISPR/Cas9 construct containing puro resistance and sgRNA targeting genes of interest. Cells were tested for cutting efficiency post selection with TIDE described earlier and by western blot. Cells were plated as spheres on growth-factor-reduced Matrigel (Corning as described previously (68) and imaged by bright field after 14 days of sphere growth.

#### Immunofluorescence

Cryosections were fixed with 4% paraformaldehyde for 10 minutes. Following fixation, slides were rinsed 3 times with PBS for 5 minutes. Samples were blocked at room temperature with blocking serum (recipe: 1% BSA, 1% gelatin, 0.25% goat serum 0.25% donkey serum, 0.3% Triton-X 100 in PBS) for 1 hour. Samples were incubated with primary antibody diluted in blocking serum overnight at 4°C followed by 3 washes for 5 minutes in PBS. Secondary antibody was diluted in blocking serum with DAPI and incubated for 1 hour at room temperature in the dark. Following incubation, samples were washed 3 times for 5 minutes in PBS. Coverslips were added on slides using MOWIOL/DABCO based mounting medium and imaged under microscope next day. For quantification, laser power and gain for each channel and antibody combination were set using secondary only control and confirmation with primary positive control and applied to all images. For paraffin sections,

#### Mammary gland whole mount

Mammary gland whole mounts were prepared as previously described for visualization of endogenous proteins and fluorescent labelling (https://doi.org/10.1016/j.ccell.2019.02.010). Briefly, 2 mm^3^ pieces of mammary gland were fixed for 45 minutes in 4% pfa, followed by a 30 minute wash in WB buffer, 2 hrs in WB1 and an overnight incubation in anti-Keratin8 and anti-Keratin14 antibodies diluted in WB2 buffer. The following day, the pieces underwent 3 x 1hr washes in WB2 buffer prior to overnight incubation in secondary antibody (at 1:200 dilution) with DAPI added at 4°C. Finally, pieces were washed 3 times for 1 hour each and then cleared using FUnGI solution for 2+ hours at room temperature until glands appeared sufficiently cleared, and then were mounted and imaged using confocal microscopy.

#### RNA-seq and GSEA analyses

Tumors were minced and treated with collagenase for 45 minutes and trypsin for 15min. Single cell suspensions from tumors were sorted to isolated GFP+ cells using fluorescence activated cell sorting (FACS). RNA was extracted from FACS-isolated cells using Quick-RNA Plus Mini Kit (Zymoresearch Inc., #R1057) as per the manufacturer’s instructions. RNA quality was assessed using an Agilent 2100 Bioanalyzer, with all samples passing the quality threshold of RNA integrity number (RIN) score of >7.5. The library was prepared using an Illumina TrueSeq mRNA sample preparation kit at the LTRI sequencing Facility, and complementary DNA was sequenced on an Illumina Nextseq platform. Sequencing reads were aligned to mouse genome (mm10) using Hisat2 version 2.1.0 and counts were obtained using featureCounts (Subread package version 1.6.3) (69).

Differential expression was performed using DESeq2 (70) release 3.8. Gene set enrichment analysis was performed using GSEA version 4.0; utilizing genesets obtained from MSigDB (https://www.gsea-msigdb.org/gsea/msigdb). For integration with human and existing mouse tumor models, clustering was conducted after normalization and filtering for only intrinsic genes as described previously (71,72). Metascape analysis was performed using default settings. g:Profiler was run using the following parameters: version e104_eg51_p15_3922dba; ordered: true; sources: GO:MF, KEGG, REAC, HPA, HP; with all other parameters at default settings.

#### Mammary gland isolation and flow cytometry for lineage tracing

Mice were injected with Ad5-K5-Cre (VVC-U of Iowa-1174) or Ad5-K8-Cre (VVC-Li-535) in the #3 or 4 mammary glands with no greater than 2 replicates of a single condition per mouse. Individual mammary glands were harvested digested according to Stemcell Technologies gentle collagenase/hyaluronidase protocol. Briefly glands we digested overnight shaking at 37°C in 250 ul Gentle Collagenase (Stemcell Technologies #07919) 2.25 ml of complete Basal Epicult media formulated according to manufacture instructions (Epicult Basal Medium Stemcell Technologies #05610, 10% Proliferation Supplement, 5% FBS, 1% Penicillin-Streptomycin, 10 ng/ml EGF, 10 ng/ml bFGF, 0.0004% heparin). Glands were then treated with ammonium chloride and triturated for 2 minutes in pre-warmed trypsin followed by dispase. Cells were stained with CD45, CD31, Ter119, CD49f and EPCAM for luminal and basal cell identification.

#### Single-cell processing and library preparation

Pik3ca^HR^ and Pik3ca^HR^ Kdm6a^KO^ were cohoused for at least 14 days prior to injection to synchronize estrus cycles, with control LSL-GFP-Cas9 mice housed separately due to limitations in mouse numbers per cage. Each mouse was injected with 5x10^8^ pfu/ml Ad-Cre or 8x10^8^ pfu/ml Ad5-Cre in the left and right #4 mammary glands. Two mice per group were harvested except for the Pik3ca^HR^Kdm6a^KO^ in the K5-Cre experiment, which was preformed on one mouse. Mammary gland digestion was carried out as described above except two glands were pooled per mouse, and glands were digested in 2x gentle collagenase/hyaluronidase for 2 hours with trituration by P1000 pipette half-way through digestion. Cells were then sorted for GFP+ infected cells and immediately processed for snATACseq or scRNAseq according to 10x Genomics protocol (scRNAseq 3’ kit v.3.1 and snATACseq kit v1.1). Approximately 5000 cells per sample were sequenced with targeted 50 000 reads/cell.

#### 10X single cell RNA-seq data processing

The raw sequencing data from each channel was first aligned in Cell Ranger 4.0.0 using a customized reference based on refdata-gex-mm10-2020-A-R26 to allow quantification of EGFP expression. The EGFP reporter transgene was added to the refdata-gex-mm10-2020-A-R26 reference and rebuilt by running cellranger mkref with default parameters (10x Genomics). To minimize the batch effects from sequencing depth variation, we further used `cellranger aggr` function to match the depth of mapped reads. The filtered gene-by-cell count matrices from 10x `cellranger aggr` were further QCed and analyzed in R package Seurat (v3.2.3) (73). Merged library was first processed in Seurat with `NormalizeData(normalization.method = “LogNormalize”)` function. The normalized data were further linear transformed by `ScaleData()` function prior to dimension reduction. Principal components analysis (PCA) was performed on the scaled data by only using the most variable 2000 genes (identified using the default “vst” method). Cell were examined in each sample across all clusters to determine the low-quality cell QC threshold that accommodates the variation between cell types. Low-quality cells were removed with the same filtering parameters on the merged object (percent.mt <=10 & nCount_RNA >= 2500 & nCount_RNA <50000 & nFeature_RNA>=1000). Stromal cell contamination from FACS-sorting and doublet clusters were removed to keep only mammary epithelium cells. QCed merged dataset was further integrated using the `RunHarmony()`function in SeuratWrappers R package to minimize the batch effect between the Ad-Cre batch and K5-Cre batch. Top 30 harmony-PCs were used for subsequent UMAP embedding and neighborhood graph construction of the integrated dataset. To investigate Ad-Cre and K5-Cre separately, the QCed dataset was split into Ad-Cre and K5-Cre subsets and then reprocessed as described above and clusters were labeled with cell types based on marker gene expression and sample/library identity. First 30 PCs in K5-Cre subset and first 40 PCs in Ad-Cre subset were selected as significant PCs for downstream UMAP embedding and neighborhood graph construction in Seurat. Pseudotime analysis was performed using Monocle3 on K5-Cre basal cells. A central point within the WT Control cluster was set as the root node and pseutotime was calculated with automatic partitioning. The ML-HS cluster was portioned separately from the remaining cells and was excluded from visualization. Diffusion mapping was performed on epithelial cells excluding the ML-HS cluster using the destiny package (v3.4.0). The first 3 eigenvectors were used for visualization using the plot3d package.

#### Cerebro shinyapp of single cell RNA-seq data

Final processed Seurat objects from the harmony integrated dataset, Ad-Cre subset, and K5-Cre subset were further processed using the cerebroApp functions in the cerebroApp R package (v1.3.0) (74). Cerebro processed data was hosted on shinyapps.io server and it is accessible though this link: https://wahl-lab-salk.shinyapps.io/Kdm6aKO_scRNAseq/.

#### 10X single nucleus ATAC-seq data processing

The raw sequencing data from each mouse was first processed separately in 10x cellranger atac 1.2.0 pipeline using refdata-cellranger-atac-mm10-1.2.0 reference. To minimize the batch effects from sequencing depth variation, we further used the `aggr` function in cellranger-atac pipeline to match the depth of mapped reads across samples. The post-normalization fragments output from the 10x cellranger-atac aggr pipeline was imported into ArchR (75) and further QCed and analyzed. Arrow files were created with the initial filtering: `minTSS=4` and `minFrags=1000`. Each library was inspected separately to determine the QC filtering thresholds. All samples were further QC filtered with `TSSEnrichment > 6` and `log10(nFrags) >= 3.4` with the exception of the WT sample, which used a higher threshold `log10(nFrags) >= 3.55`). The merged samples were first embedded in UMAP by first running latent semantic indexing with 1 iteration with the `interativeLSI` function. Clusters identify were inferred based on the gene score of marker genes. Clusters of doublets, are marked by shared marker gene expression from two different lineages and higher number of reads per cell on average as previously described (23). Clusters of stromal cell contamination and doublets were removed from subsequent analysis based on marge gene expression and average read-depth distribution as previously described (23). The cleaned mammary epithelial cell dataset was re-processed through a 1-iteration `interativeLSI` with default parameters. All top 20PCs were used to embed cells in two dimensional UMAP. Clusters were called by using the `addClusters(method=’Seurat’,resolution=1.1,dimsToUse=1:20)` function and subsequently labelled with cell types using gene scores of marker genes and sample identity. Pseudo-bulk profile with replicates was generated and reproducible peaks were identified by calling peaks specifically in each clusters or cell types across replicates using the macs2 method. Differentially accessible peaks were identified using the `getMarkerFeatures` and `getMarkers (cutOff = “FDR <= 0.1 & Log2FC >= 1”)`. TF motif activity was inferred by using the chromVAR TF enrichment deviation z-scores in ArchR (24,75). Heatmaps were generated using the ComplexHeatmap R package using scaled and centered values across cell type groups (76).

#### TCGA data

Clinical and pathological data, somatic genetic mutations and genomic copy numbers were obtained from the cBioPortal (77). Gene expression (RNA-seq fragments per kilobase of transcript per million mapped reads (FPKM) upper quartile normalized (UQ)) data were obtained from the Genomic Data Commons Data Portal (https://portal.gdc.cancer.gov). In survival analyses, EpiDriver mutations were defined as somatic gene mutations and/or homozygous genomic deletions of *ASXL2*, *BAP1*, *KMT2C*, *KMT2D*, *KDM6A*, and/or *SETD2*. The TCGA breast cancers were previously scored for PI3K/AKT/mTOR signaling using a transcription-based CMAP signature, in which high values were associated with poor outcome (78). The measures of phospho-Ser473 AKT were downloaded from The Cancer Proteome Atlas (TCPA) (36) and corresponded to level 4 normalized values from assays using reverse-phase protein arrays. The high/low threshold (value = 0) of CMAP and pAKT were confirmed by examining the value distributions in all primary tumors. The Kaplan-Meier curve and log-rank test analyses were performed in R software using the *survival* and *survminer* packages.

#### Statistics and reproducibility

All quantitative data are expressed as the mean ± SE. Differences between groups were calculated by two-tailed Student’s t-test, Wilcoxon Rank-Sum test (when data was not normally distributed) or Log-rank test for survival data using Prism 7 (GraphPad software). Where adjustment is indicated and the method not otherwise specified, p value was adjusted using Bonferroni correction.

## Supplementary Figure legends

**Supplementary Fig. S1. Direct *in vivo* CRISPR knock-out efficiency in mouse mammary epithelium.**

**a,** Experimental schematic showing genetic ablation of green fluorescent protein (GFP) from double reporter R26-LSL-Cas9; GFP^het^; R26-LSL-tdTomato^het^ mice transduced with an sgGFP-Cre lentivirus that targets GFP or non-targeting control lentivirus (sgNT). **b**, Representative images showing GFP and tdTomato expression in R26-LSL-Cas9-GFP^het^; R26-LSL-tdTomato^het^ mammary epithelium transduced with sgGFP-Cre lentivirus or control non-targeting sgNT-Cre lentivirus. **c,** Flow cytometry analysis shows the percent of GFP+/tdTomato+ double positive or tdTomato+ single positive cells. **d**, Representative images and FACS plots depicting red fluorescence in the mammary epithelium of *Urod* mosaic knock-out mice indicative of efficient, bi-allelic mutagenesis of *UroD* gene. **e**, Tumor-free survival for *Pik3ca*^H1047R^;Cas9 mice transduced with lentiviral sgRNAs targeting the *Trp53* gene or the Tigre ‘safe harbor’ locus as control. (n≥5 per group; p<0.0001). **f**, Representative images of mammary epithelium transduced with a Lenti-GFP/Lenti-RFP mixture showing cells transduced with GFP or RFP or double-infected expressing GFP+/RFP+ cells. More double positive cells are observed at higher viral titre. **g**, Whole mount image of a mammary gland transduced with Lenti-Cre in a R26-LSL-tdTomato^het^ mouse. **h and i,** Representative FACS plot (h) and quantification (i) for RFP+ cells in Lenti-Cre injected mammary glands from R26-LSL-tdTomato^het^ mice.

**Supplementary Fig. S2. Library screening and hit validation. a,** Graph showing sgRNA representation from breast cancer libraries in plasmid DNA versus DNA from infected MEFs. Each dot represents a guide. Full representation is maintained with high correlation in abundance. **b,** Representative pie charts showing tumor suppressor genes with enriched sgRNAs in tumor DNA obtained from three different mammary gland tumors. **c,** METASCAPE pathway enrichment analysis within screen hits. **d,** Tumor-free survival of Pik3ca^H1047R^ transduced individually with sgRNAs targeting the indicated genes. sgRNAs from the original library screen (‘Lib’) or independently designed sgRNAs (‘New’) were used. No significant survival difference was found between pairs of sgRNAs targeting the same gene. **e,** Tumor-free survival of Pik3ca^H1047R^ transduced individually with sgRNAs targeting Bap1 or the inert Tigre locus as control. **f,** Gene editing efficiency was determined using sanger-sequencing data of PCR-amplified sgRNA target sites followed by Tracking of Indels by Decomposition (TIDE https://tide.nki.nl) algorithm on bulk or GFP+ sorted tumor cells. Colour denotes sgRNA as shown in (c). **g,** Representative sanger sequencing plots showing discordance of DNA from a sgAsxl2-targeted sample compared to a wildtype sample. **h-k,** Western blot analysis showing loss of protein expression after CRISPR-mediated knockout of Apc (h), p53 (i), Asxl2 (j) and Kdm6a (k). **l,** Western blot analysis of Kdm6a in wild-type, Kdm6a^fl/+^ heterozygous and Kdm6a^fl/fl^ homozygous mammary tumors with densitometry of Kdm6a normalized to loading control H3 (**m**).

**Supplementary Fig. S3. Histological characterization of CRISPR knockout tumors. a,** Pathological characterization of mammary tumors from each knockout. **b,** Representative H&E from sgAPC and sgSetd2 tumors showing adenosquamous carcinoma (top) and complex adenocarcinoma (bottom). **c and d,** Quantification of estrogen receptor staining intensity in the nucleus (c) or cytoplasm (d) determined by anti-ERα immunofluorescent staining. **e and f,** Representative images of K8 and K14 staining in sgTp53 and sgNT tumors (e) and in a sgKdm6a tumor (f). **g,** Quantification of K8 and K14 staining in tumors transduced with the indicated sgRNAs.

**Supplementary Fig. S4. Transcriptional profiling of epigenetic knockout tumors**. **a,** Dot plot showing GSEA for hallmark pathways in knockout tumors relative to control tumors. Circle size denotes -log(FDR) with lower cutoff of -log(FDR)=0.6 (FDR=0.25). Color depicts Normalized Enrichment Score (NES) with red denoting enriched pathways and blue denoting depleted pathways. **b,** GSEA plot showing enrichment of ‘GO_keratinization’ in sgApc tumors compared to control sgNT tumors. **c and d,** GSEA plots showing enrichment of ‘HALLMARK_EPITHELIAL_MESENCHYMAL_TRANSITION’(**c**) and depletion of ‘GO_EPITHELIAL_ CELL_DIFFERENTIATION’ (**d)** in EpiDriver knockout vs control sgNT tumors. **e,** Cross- and interspecies unsupervised clustering of CRISPR/Cas9 EpiDriver knockout Pik3ca^H1047R^ tumors as well as Ad-K5-Cre Pik3ca^H1047R^ and Ad-K5-Cre Pik3ca^H1047R^;Kdm6a^fl/fl^ tumors compared to previously described mouse breast cancer models and human breast cancer subtypes.

**Supplementary Fig. S5. scRNAseq reveals lactation mimicry. a,** Contribution the genotypes to each cell clusters. **b,** Unsupervised UMAP plot of scRNAseq profile colored by genotype and UMAP and violin plots depicting scRNAseq data of selected alveolar/lactation markers. Single gene expression on UMAP plots is shown with a maximum cutoff of q90.

**Supplementary Fig. S6. scRNAseq reveals lineage plasticity. a,** UMAP and violin plot showing EMT and hypoxia signatures. **b and c**, UMAP and violin plots depicting scRNAseq data of selected HS-ML (b) and basal (c) markers. Single gene expression on UMAP plots is shown with a maximum cutoff of q90.

**Supplementary Fig. S7. snATACseq reveals significant changes in chromatin accessibility upon Kdm6a loss. a,** Unsupervised clustering of chromVAR TF activity scores for 788 transcription factors in the major clusters obtained by snATACseq UMAP clustering. **b,** UMAP blot of snATACseq profile colored by genotype and clusters (top) and UMAP plots depicting Ehf (middle) and Elf5 (bottom) ChromVAR TF activity score.

**Supplementary Fig. S8. snATACseq confirms lactation mimicry and reveals WNT as well as NOTCH signaling in BA2 cluster. a,** Unsupervised clustering of ArchR gene accessibility score across the major clusters obtained by snATACseq. **b and c**, METASCAPE pathway enrichment analysis showing enriched pathways shared between BA2 and LP2 clusters compared to all other clusters (b) and between BA2 compared to the basal clusters (c) as inferred by gene accessibility scores obtained by snATACseq. **d and e,** UMAP and violin plot showing WNT protein binding and signaling (d) and NOTCH regulation and signaling (e) signatures by scRNAseq.

**Supplementary Fig. S9. BA2 cluster exhibits downregulated basal markers. a,** Unsupervised UMAP blot of snATACseq profile showing single-cell gene accessibility scores of the indicated basal cell markers revealing downregulation in the BA2 cluster. **b and c**, Signal tracks of the aggregate snATAC profile for Krt5 (b) and Trp63 (c) and the indicated cluster.

**Supplementary Fig. S10. LP2 cluster exhibits downregulated luminal markers. a,** Unsupervised UMAP blot of snATACseq profile showing single-cell gene accessibility scores of the indicated luminal cell markers revealing downregulation in the LA2 cluster. **b and c**, Signal tracks of the aggregate snATAC profile for Ehf (b) and Elf5 (c) and the indicated cluster.

**Supplementary Fig. S11. Lineage tracing of basal and luminal cells. a and b,** Schematic (**a**) and representative immunofluorescent whole mount image (**b**) illustrating the location of basal K5+/K14+ and luminal K8+/K18+ mammary epithelial cells. **c,** Representative FACS blot depicting lineage tracing strategy and lineage restriction of Ad-K5-Cre and Ad-K8-Cre viral particles. **d,** Percent of GFP+ EPCAM^hi^CD49f^mid^ luminal cells at different time points after Ad-K5-Cre injection into mammary epithelium of R26-LSL-GFP mice as determined by flow cytometry. **e,** Percent of GFP+ EPCAM^med^CD49f^hi^ basal cells at different time points after Ad-K8-Cre injection into mammary epithelium of R26-LSL-GFP mice as determined by flow cytometry. **f,** Percent of GFP+ EPCAM^hi^CD49f^mid^ luminal cells at 5.5 weeks after Ad-K5-Cre injection into mammary epithelium of heterozygous Kdm6a^fl/+^;Pik3ca^H1047R^;R26-LSL-GFP mice as determined by flow cytometry. **g,** Percent of GFP+ EPCAM^mid^CD49f^hi^ basal cells at different time points after Ad-K8-Cre injection into mammary epithelium of Pik3ca^H1047R^, Pik3ca^H1047R^;Kdm6a^fl/fl^ and Pik3ca^H1047R^;Asxl2^fl/fl^ mice.

**Supplementary Fig. S12. scRNAseq reveals transdifferentation of basal to alveolar cells. a,** Contribution the genotypes to each cell clusters. **b-e,** UMAP and violin plots of K5-Cre scRNAseq data of the indicated basal cell markers (b) and luminal progenitor markers (c).

**Supplementary Fig. S13. scRNAseq reveals transdifferentation of basal to alveolar cells. a-c,** UMAP and violin plots of K5-Cre scRNAseq data of the indicated lactation markers (a) and HS-ML markers (b). **d,** Plot showing expression of alveolar marker Wfdc18 and Acta2 in scRNAseq data. Of note, many Pik3ca;Kdm6a^KO^ express both markers.

**Supplementary Fig. S14. scRNAseq reveals heterogeneity along Pik3ca^H1047R^;Kdm6a^KO^ cells. a,** METASCAPE pathway enrichment analysis showing enriched pathways in Kdm6a_M versus Kdm6a_L clusters. **b,** Diffusion mapping of scRNAseq data displayed by a 3D plot of the first 3 eigenvectors depicting branching of Kdm6a^KO^ cells between Kdm6a_B and Kdm6a_L clusters. **c,** Diffusion mapping of scRNAseq data displayed by a 3D plot of the first 3 eigenvectors depicting alveolar marker *Csn3* and the basal marker *Igfbp2* showing separation of the Kdm6a clusters. **d,** UMAP and violin plot of scRNAseq data showing expression of the genes relatively specific to the Kdm6a_L cluster.

**Supplementary Fig. S15. Integration of Ad-Cre and Ad-K5Cre scRNAseq. a,** UMAP plots of scRNAseq data of Ad-Cre and Ad-K5Cre scRNAseq colored by genotype (left), by epithelial lineage (middle) and identified clusters (right). **b,** UMAP plots of the combined scRNAseq data showing only the Ad-Cre data (left), only the Ad-K5-Cre data (middle) and with Ad-K5-Cre Pik3ca;Kdm6a^KO^ highlighted in red (right). **b,** UMAP plots of the combined scRNAseq data showing only the Ad-Cre data showing basal (top left), alveolar/lactation (top right), luminal progenitor (bottom left), and involution (bottom right) marker signatures.

**Supplementary Fig. S16. scRNAseq reveals Rb1/E2F pathway in basal proliferating cells and marker genes of BA-alveolar-LP transdifferentation. a,** UMAP and violin plots of scRNAseq data showing expression of the proliferation marker Mki67 (KI67) and the indicated E2F/Rb1 target gene.

**Supplementary Fig. S17. EpiDriver loss triggers transformation of MCF10A-PIK3CA^H1047R^ cells**

**a and b,** CRISPR/Cas9-mediated knock-out efficacy in MCF10A-PIK3CA^H1047R^ cells transduced with Cas9-Puro-sgRNA lentivirus targeting the indicated gene. TIDE analysis showing indel abundance (a) and western blot showing protein levels (b). **c,** MCF10A^H1047R^ cells with the indicated gene knockouts grown as spheres in matrigel without EGF. **d**, Tumor histology for sgKDM6A MCF10A-PIK3CA^H1047R^ tumor. Inset shows invasion into surrounding tissue. Scale bar = 50μm.

**Supplementary Fig. S18. Epigenetic regulators are mutated in human tumors and suppress transformation of MCF10A cells. a,** Oncoprint of the indicated epigenetic genes identified as tumor suppressors in our screen compared to *KMT2A/B*, *MEN1* and COMPASS-like core proteins (ASH2L, DPY30, RBBP5 and WDR5) as well as alterations in *PIK3CA*. Shallow deletion shown only for KDM6A. Interestingly, KMT2A/B methyltransferases and their interaction partner *MEN1* or genes coding core COMPASS-like complex proteins common to all COMPASS complexes are not mutated in breast cancer. **b,** *KDM6A* mRNA expression stratified by copy-number alteration. Data for **a,** and **b,** were obtained from METABRIC and accessed through cBioPortal. **c,** DSS of breast cancer patients in the TCGA cohort stratified by PI3K CMAP transcriptional signature and EpiDriver mutations. The number of patients at risk and long-rank *p* value are shown. **d,** DSS stratified by tumor subtype, and PI3K CMAP transcriptional signature and EpiDriver mutations. There was a limited set of events in luminal A tumors to evaluate this association, but this subtype showed a significant higher proportion of tumors with high PI3K and EpiDriver mutation (odds ratio = 12.2; *p* < 2x10-16). **e,** DSS stratified by tumor subtype, and phospho-Ser473 AKT and EpiDriver mutations.

